# DDX5 targets tissue-specific RNAs to promote intestine tumorigenesis

**DOI:** 10.1101/2020.03.25.006668

**Authors:** Nazia Abbasi, Tianyun Long, Yuxin Li, Evelyn Ma, Brian A. Yee, Parth R. Patel, Ibrahim M Sayed, Nissi Varki, Soumita Das, Pradipta Ghosh, Gene W. Yeo, Wendy J.M. Huang

**Author notes:** These authors contributed equally (Nazia Abbasi, Tianyun Long).

## Abstract

Tumorigenesis in different segments of the intestinal tract involves tissue-specific oncogenic drivers. In the colon, complement component 3 (C3) activation is a major contributor to inflammation and malignancies. By contrast, tumorigenesis in the small intestine involves fatty acid-binding protein 1 (FABP1). However, little is known of the upstream mechanisms driving their expressions in different segments of the intestinal tract. Here, we report that an RNA binding protein DDX5 augments C3 and FABP1 expressions post-transcriptionally to promote tumorigenesis in the colon and small intestine, respectively. Mice with epithelial-specific knockout of DDX5 are protected from intestine tumorigenesis. The identification of DDX5 as the common upstream regulator of tissue-specific oncogenic molecules provides a new therapeutic target for intestine cancers.

## Introduction

Tissue-specific oncogenic molecules drive tumorigenesis in different segments of the intestine tract. In the colon, complement component C3 protein induces the expression of pro-inflammatory cytokines, such as IL-1b and IL-17 [1-3]. Ablation of C3 genetically protects against tumorigenesis in high-fat diet and colitis-associated colorectal cancer models [4-7]. In the small intestine, fatty acid-binding protein 1 (FABP1) is critical for intestinal absorption of dietary long-chain fatty acids [8-10]. Intriguingly, ablation of FABP1 genetically protects against tumorigenesis in the small intestine [11].

Regulations of C3 and FABP1 expression at the transcriptional level are well described. *C3* transcription is induced by pro-inflammatory cytokines, such as TNFα, IFNγ, and IL1β [12-14], under the regulation of critical transcription factors, including the twist basic helix– loop–helix transcription factor 1 (TWIST1), CCAAT/enhancer-binding protein β (C/EBPβ), nuclear receptors Farnesoid X Receptor (FXR), and Peroxisome Proliferator-Activated Receptor α (PPARα) [15-18]. The transcription of *Fabp1* is controlled by GATA Binding Protein 4(GATA4), C/EBP, PPARα, Pancreatic And Duodenal Homeobox 1 (PDX1), and Hypoxia-Inducible Factor (HIF1α) [19-23]. However, little is known about how C3 and FABP1 expression are regulated post-transcriptionally in intestinal epithelial cells.

Post-transcription regulation of gene products is facilitated by a large number of RNA-binding proteins [24]. One member of the DEAD-box containing RNA binding protein family, DDX5, is abundantly expressed in the intestine epithelium [25]. Mutation and overexpression of DDX5 are often found in human cancers and its dysregulated expression predicts advanced clinical stage and poor survival in colorectal cancer patients [26-28]. In a xenograft model, DDX5 is responsible for the hyperproliferation of gastric cancer cells [29].

Mechanistically, DDX proteins have two major modes of action. First, they can directly bind to specific RNA substrates and utilize ATP hydrolysis energy to unwind RNA duplexes, facilitate RNA annealing, organize RNA-protein complex assembly [30], and/or promote post-transcriptional processing [31-33]. Second, DDXs can partner with transcription factors to modulate gene transcription [25, 30, 34-40]. In human cancer cell lines, DDX5 interacts with β-catenin protein and a long non-coding RNA NEAT1 to promote oncogene expressions [41, 42]. However, we know little about how the RNA binding properties of DDX5 contributes to shaping the epithelial RNA regulome during homeostasis and tumorigenesis *in vivo*.

Here, we revealed that DDX5 binds to *C3* and *Fabp1* mRNA and promotes their expressions in primary intestinal epithelial cells from the colon and small intestine, respectively. Loss of DDX5 expression in the intestinal epithelial cells protects against colonic and small intestine tumorigenesis *in vivo*. Identification of DDX5 as a common upstream regulator of tissue-specific oncogenic molecules in the intestine provides a excellent therapeutic target for treating intestinal cancers.

## Results

### DDX5 regulates RNA programs in colonic epithelial cells

In the intestinal epithelial cells (IECs) isolated from the colon and small intestine of adult wild-type (WT) mice, 35 RNA-binding DEAD-box containing proteins (DDXs) were expressed at various levels (Fig. S1A-1B). Among these, *Ddx5* was the most abundant (Fig. 1A). Western blot analyses confirmed that DDX5 proteins were present throughout the intestinal tract, and its highest expression were found in the duodenum and colon (Fig. 1B). Immunohistochemistry (IHC) and nuclear-cytoplasmic fraction revealed that DDX5 proteins predominantly localized to the nucleus of colonic IECs (Fig. 1C and S1C). Given the nuclear localization of DDX5, we hypothesize that it may bind to target IEC RNAs in the nucleus and regulate their expressions post-transcriptionally.

**Figure 1.**
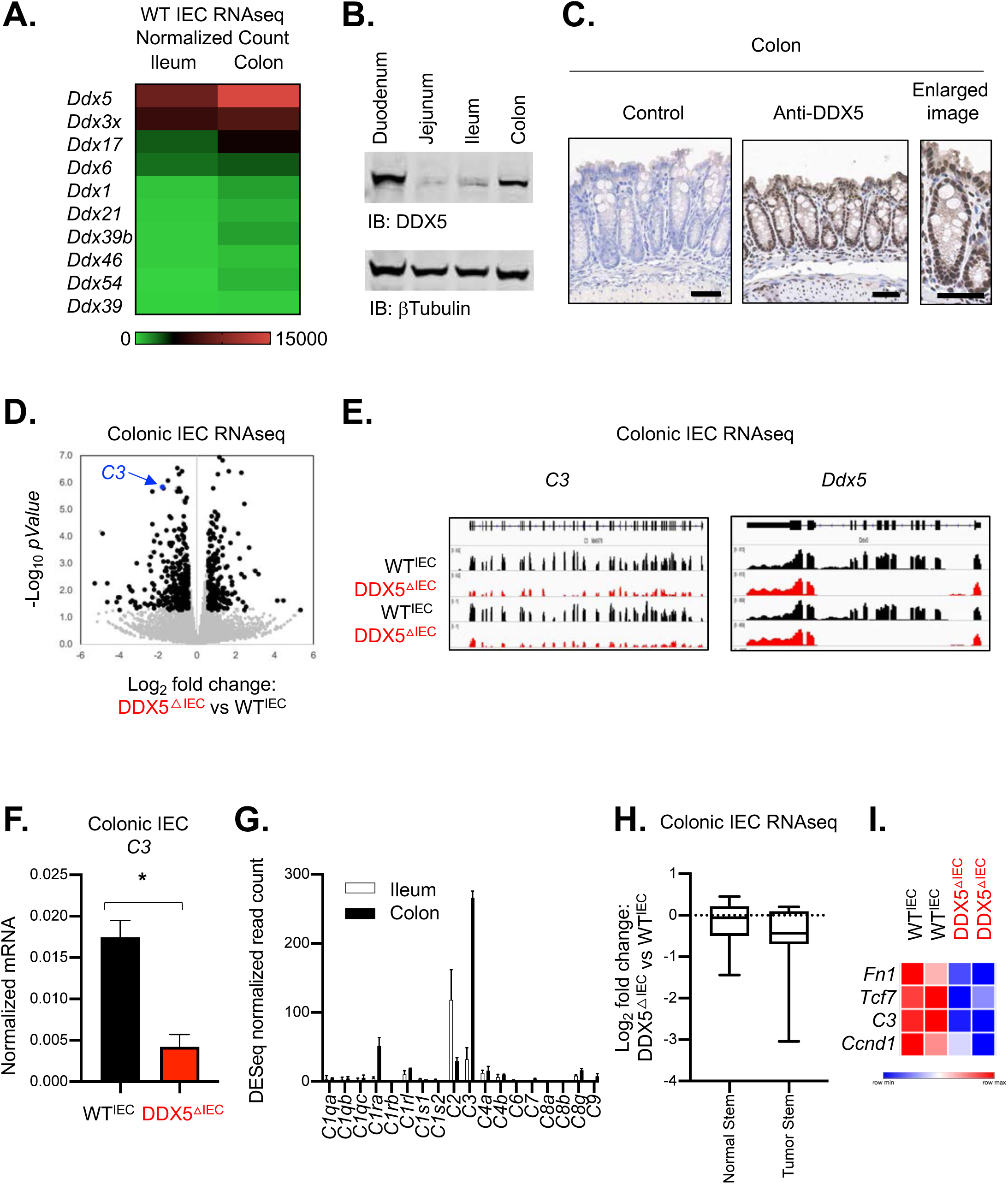
DDX5 regulates colonic epithelial RNA programs. A. Heatmap of normalized RNAseq read counts of the 10 highest expressed members of the DDX family in ileum and colon of steady-state WT mice. B. Representative western blots showing DDX5 and βTubulin protein expression in IECs from different sections of the intestine in WT mice. Experiments were repeated three times using independent biological samples with similar results. C. Representative images from IHC analysis of DDX5 in the colon of WT mice. Enlarged image is shown on the right. Scale bar represents 50 μm. D. Scatterplot of log_2_(fold changes: DDX5^ΔIEC^ over WT^IEC^) and −log_10_(*p*-Values) of colonic IEC transcripts. Black dot: all DDX5-dependent transcripts. Blue dot: *C3*. E. IGV browser displaying RNA expression at the *C3* and *Ddx5* locus in colonic IECs from two independent pairs of WT^IEC^ and DDX5^ΔIEC^ littermates. F. qPCR validation of colonic *C3* expression in 3 additional independent pairs of WT^IEC^ and DDX5^ΔIEC^ animals. Data shown are means ± standard deviations. * p<0.05 (*t*-test). G. Normalized RNAseq read counts of the complement gene family in ileal and colonic IECs from steady-state WT mice. H. Log_2_ RNA expressions fold changes of normal and tumor stem cell signatures in colonic IECs from DDX5^ΔIEC^ and WT^IEC^ littermates. I. Heatmap displaying RNA abundance of selected differentially expressed tumor stem cell transcripts in two biologically independent pairs of WT^IEC^ and DDX5^IEC^ mice.

Hence, we generated an epithelial DDX5 knockout mice (DDX5^ΔIEC^) using the Villin1 (*Vil1*)-Cre recombination system (Fig. S2A). WT^IEC^ and DDX5^ΔIEC^ mice were born in Mendelian ratios and had similar growth curves (Fig. S2B). IECs isolated from different segments of the intestinal tract confirmed efficient knockout of DDX5 at the RNA and protein levels throughout the small intestine and colon (Fig. S2C-D). Comparison of the RNA profile of colonic IECs isolated from steady-state WT^IEC^ and DDX5^ΔIEC^ mice revealed that knocking out DDX5 resulted in a downregulation of 306 and upregulation of 174 colonic IEC transcripts (Fig. 1D). Surprisingly, unlike previous cell culture studies [42], the DDX5-dependent transcripts we identified in colonic IECs were not enriched in cell proliferation, cell cycle, or apoptosis (Fig. S3A). Instead, DDX5-dependent RNA programs of the colonic IECs were enriched with genes involved in immune responses, including the completement protein-encoding transcript *C3* (Fig. 1E-F).

### Epithelial DDX5 promotes colonic tumorigenesis

The expression of the complement family of genes is tissue-specific. While the small intestine IECs predominantly expressed *C2*, colonic IECs express a high level of *C3* (Fig. 1G). The completement proteins drives pro-inflammatory cytokines expression and can activate the Wnt/β-catenin signaling cascade [1-3, 43, 44]. Numerous targets downstream of Wnt/β-catenin are well-known oncogenic drivers expressed in tumor stem cells [45]. In the C3 knockout mice, tumorigenesis in the colon is significantly blunted [6, 46]. Two independent studies in human colorectal cancer patients revealed that higher expression of *C3* predicts poor overall and relapse-free survival (Fig. S4A) [47, 48]. Therefore, we hypothesize that the loss of colonic *C3* expression would limit the expression of tumor stem cell gene program, and thus reducing the risk for colonic tumors development in the DDX5^ΔIEC^ mice. Indeed, several tumor stem cell signature genes were downregulated in DDX5^ΔIEC^ colonic IECs, including *Ccnd1, Fn1*, and *Tcf7* (Fig. 1H-I and S4B).

To test our hypothesis *in vivo*, we crossed the DDX5^flox^ line to the adenomatous polyposis coli mutant mice (APC^fl/+^CDX2Cre^+^, aka *Apc*^ΔcIEC^) (Fig. 2A). APC mutant mice are used to model familial adenomatous polyposis in human, where haploinsufficiency of the tumor suppressor APC leads to aberrant β-catenin activation and the development of large colonic adenomas that progresses to invasive carcinomas [49, 50]. In this model, the loss of one copy of the *Apc* allele in the epithelium results in spontaneous weight loss and tumor lesions as early as three-months of age [51-53]. Periodic Acid-Schiff (PAS) stained histological sections of colonic adenomas from the *Apc*^ΔcIEC^ mice showed a loss of differentiated goblet cell population and neoplastic cell infiltration beyond the basal membranes (Fig. S5A). Western blot analyses revealed that DDX5 proteins were expressed at a significantly higher level in colonic tumors from *Apc*^ΔcIEC^ mice compared to adjacent normal tissues or IECs from non-tumor bearing WT mice (Fig. S5B), similar to findings previously reported in human CRCs [26]. At 4-months of age, *Apc*^ΔcIEC^ *Ddx5*^ΔcIEC^ mice had lower incidence of anal prolapse (Fig. 2B) and experienced less weight loss compared to *Apc*^ΔcIEC^*Ddx5*^WT^ controls (Fig. 2C). In the colons harvested, *Apc*^ΔcIEC^*Ddx5*^ΔcIEC^ mice had fewer macroscopic tumors (Fig. 2D-E). IHC studies confirmed that DDX5 was indeed knocked out and lesions from the *Apc*^ΔcIEC^*Ddx5*^ΔcIEC^ mice had reduced expression of the cell proliferation marker, Ki67 (Fig. S5C). Notably, the phenotype of the *Apc*^ΔcIEC^ *Ddx5*^ΔcIEC^ mice mirrored those observed in the C3-deficient animals [4-7]. Loss of APC causes adenomas, but not adenocarcinomas [54]. To assess the role of DDX5 in colorectal cancers, relapse-free survival in CRC patients was examined. Intriguingly, a decrease of relapse-free survival correlates with higher expression of DDX5 (Fig. 2F). Together, these *in vivo* results demonstrate DDX5 as a potent driver of colonic tumorigenesis in both mice and human.

**Figure 2.**
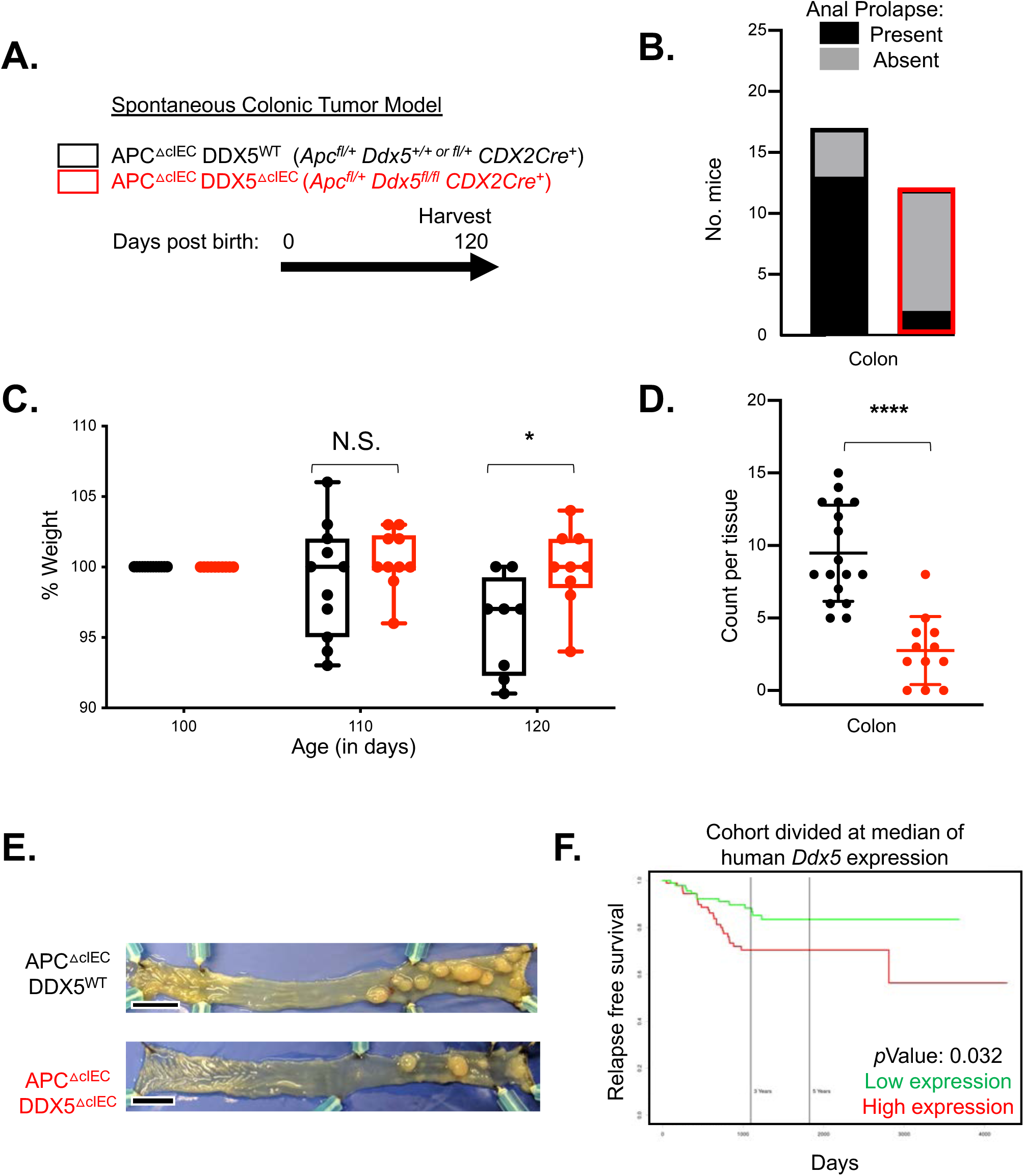
DDX5 promotes colonic tumorigenesis in *Apc* mutant mice. A. Tumor bearing APC^ΔcIEC^DDX5^WT^ (*n*=12) and APC^ΔcIEC^ DDX5^ΔcIEC^ (*n*=13) littermates. B. Anal prolapse incidents recorded in mice described in A. C. Percent weight change of each mice in A on day 110 and 120 compared to day 100. Each dot represents one mouse. Data shown are means ± standard deviations. * p<0.05 (*t*-test). D. Colonic tumor counts from APC^ΔcIEC^DDX5^WT^ (*n*=12) and APC^ΔcIEC^ DDX5^ΔcIEC^ (*n*=13) tumor-bearing animals. Each dot represents one mouse. Data shown are means ± standard deviations. **** *p-value* <0.0001 (*t*-test). E. Representative bright-field images of tumor-bearing colons from APC^ΔcIEC^DDX5^WT^ and APC^ΔcIEC^ DDX5^ΔIEC^ animals. F. PROGgeneV2 view of relapse-free survival in 187 CRC patient cohort (GSE14333) divided at median of human *Ddx5* gene expression.

### Epithelial DDX5 directly binds *C3* RNA and enhances its expression post-transcriptionally

Next, we asked whether DDX5 regulation of *C3* is intrinsic to colonic epithelial cells and independent of inputs from gut microbiota and immune cells using an organoid culture system. Briefly, colonic crypts containing epithelial stems cells were harvested from WT^IEC^ and DDX5^ΔIEC^ littermates and maintained *ex vivo* for 5-7 passages in the presence of EGF, Noggin, and R-spondin (Fig. 3A). RNAseq of these colonic organoids revealed a similar reduction of *C3* RNA in DDX5-deficient cultured cells (Fig. 3B), suggesting that the regulation of *C3* by DDX5 is epithelial cell-intrinsic.

**Figure 3.**
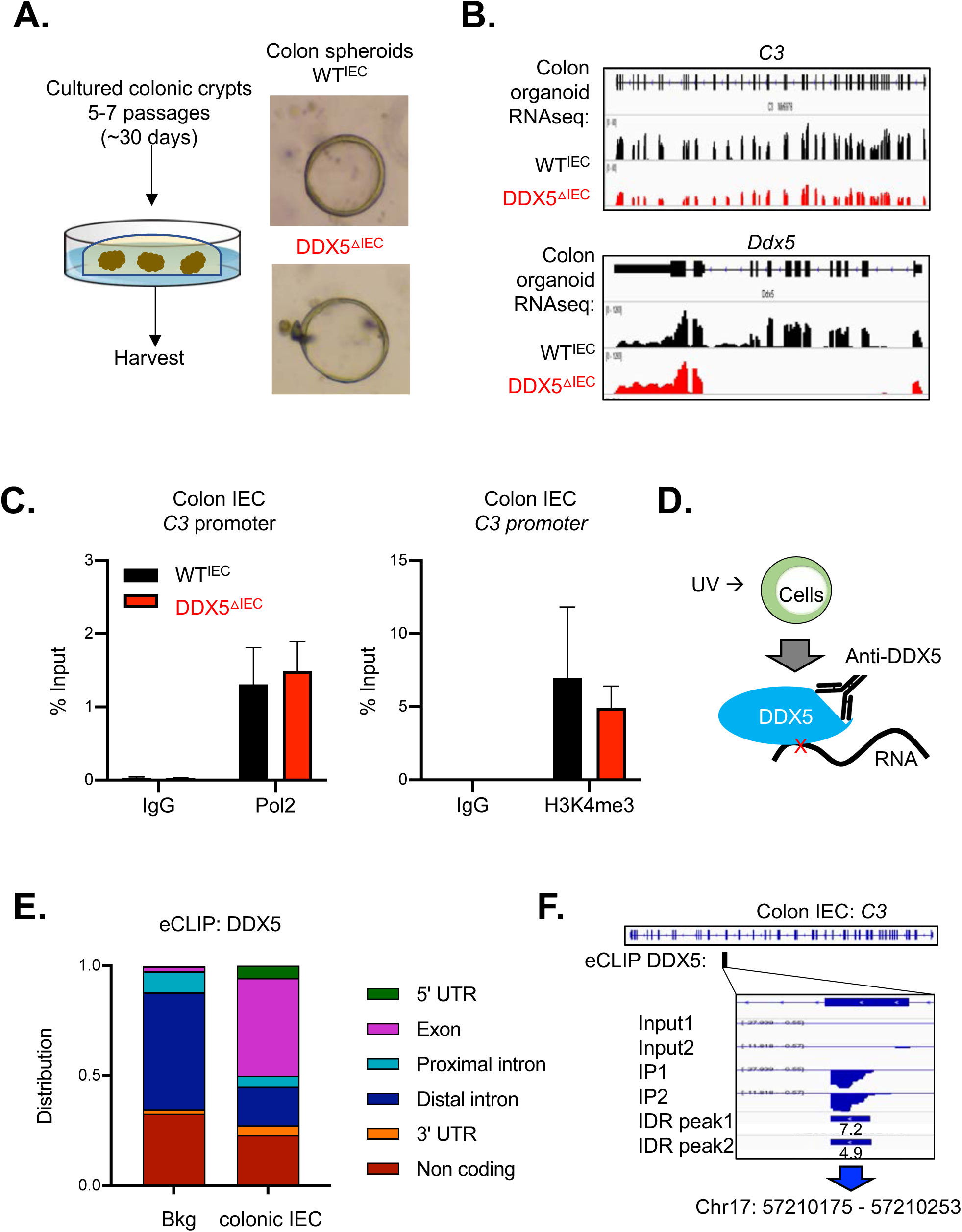
Epithelial DDX5 binds *C3* RNA to enhance its expression post transcriptionally. A. Representative bright field images of organoids cultured from colonic crypts of WT^IEC^ and DDX5-deficient mice following the protocol outlined in [66]. B. IGV browser displaying RNA expression at the *C3* and *Ddx5* locus in cultured organoids derived from WT^IEC^ and DDX5^ΔIEC^ colonic crypts. C. Chromatin immunoprecipitation (ChIP-qPCR) assay of RNA polymerase II and H3K4me3 in colonic IECs from 2 independent pairs WT^IEC^ and DDX5-deficient mice. D. Schematic representation of the enhanced CLIP (eCLIPseq) experiment. WT IECs were subjected to UV-mediated crosslinking, followed by lysis, treatment with RNases, and immunoprecipitation (IP) of protein-RNA complexes using an anti-DDX5 antibody (see Table S1). RNA fragments protected from RNase digestion were subjected to linker ligation and reverse-transcription to generate eCLIPseq libraries for high-throughput sequencing on the Illumina platform. E. DDX5 binding preference as identified by eCLIPseq on the different colonic RNA regions. Background (Bkg) is defined as the RNA regions in the annotated mouse transcriptome from GENCODE. F. IGV browser displaying the DDX5 binding to *C3* RNAs in WT colonic IECs as defined by eCLIPseq.

If DDX5 promotes *C3* expression at the transcription level, we expect to observe altered RNA polymerase II recruitment and H3K4me3 deposition on the *C3* gene promoter in colonic IECs from DDX5^ΔIEC^ mice. However, chromatin Immunoprecipitation (ChIP) qPCR assay showed that a similar enrichment of RNA polymerase II and H3K4me3 were found on the *C3* promoter in colonic IECs from WT^IEC^ and DDX5^ΔIEC^ mice (Fig. 3C). Therefore, we hypothesize that DDX5 may bind to and regulate *C3* transcripts at the post-transcriptional level. To test this, we performed the enhanced cross-linked immunoprecipitation (eCLIPseq) assay on two biological replicates of UV crosslinked colonic IECs from WT mice using the anti-DDX5 antibodies (Fig. 3D and Table S1). Successful pull-down of DDX5 proteins were confirmed by western assays (Fig. S6A). Sequencing results were processed by the ENCODE eCLIPseq analysis pipeline, as described in [55] and outlined in Fig. S6B. Using a cutoff of 3 in both log_10_ *p*-values and log_2_ fold changes of IP signal over input, we identified 201 colonic IEC RNA sites, corresponding to 139 transcripts, that were significantly enriched by the anti-DDX5 antibodies. Over 44% of the DDX5-bound sites on colonic IEC RNAs localized to coding regions (Fig. 3E).

Only three of the 139 DDX5-bound transcripts, including *C3*, experienced significant RNA expression changes in DDX5-deficient colonic IECs. On the *C3* RNA, DDX5 was enriched on a region encoded by exon 30 (Fig. 3F).

*C3* RNAs from mouse and human have over 80% conservation in identity. Therefore, we speculated that the DDX5 regulation of *C3* RNA we found in mouse may be conserved in human colonic epithelial cells. Indeed, RNAi mediated knockdown of human *Ddx5* in Caco-2 cells where transcription was inhibited by flavipiridol resulted in a reduction of h*C3* RNA levels (Fig. S7A-B). Furthermore, insertion of the short stretch of DDX5 bound region of mouse *C3* into the 3’UTR of the psiCheck2 reporter was sufficient to potentiate *Renilla* luciferase activities in a DDX5-dependent manner in another human epithelial cell line (SW480) (Fig. S7C). Together, these results highlight an epithelial intrinsic, direct, and conserved role of DDX5 in post-transcriptionally promoting *C3* expression.

### DDX5 binds a distinct set of RNAs to drive tumorigenesis in the small intestine

Similar to the colon, DDX5 is abundantly expressed in the small intestine IECs under steady state (Fig. 1A and 4A). We asked whether DDX5 also promote tumorigenesis in the small intestine. Epithelial DDX5 knockout mice (DDX5^flox^Vil1Cre^+^) were crossed to the adenomatous polyposis coli (APC) mutant mice (APC^fl/+^) (Fig. 4B). While the APC^fl/+^CDX2Cre^+^ mutant mice only develop colonic tumors (Fig. 2), *Apc*^ΔIEC^ mice harbor intestinal tumors in both the small intestine and colon. By 110 days of age, *Apc*^ΔIEC^*Ddx5*^ΔIEC^ littermates continued to gain weight, but *Apc*^ΔIEC^ DDX5^WT^ mice began to experience significant weight loss (Fig. 4C). On day 120, we found fewer macroscopic lesions in the colon and small intestine of *Apc*^ΔIEC^*Ddx5*^ΔIEC^ mice compared to their littermate controls (Fig. 4D-E).

**Figure 4.**
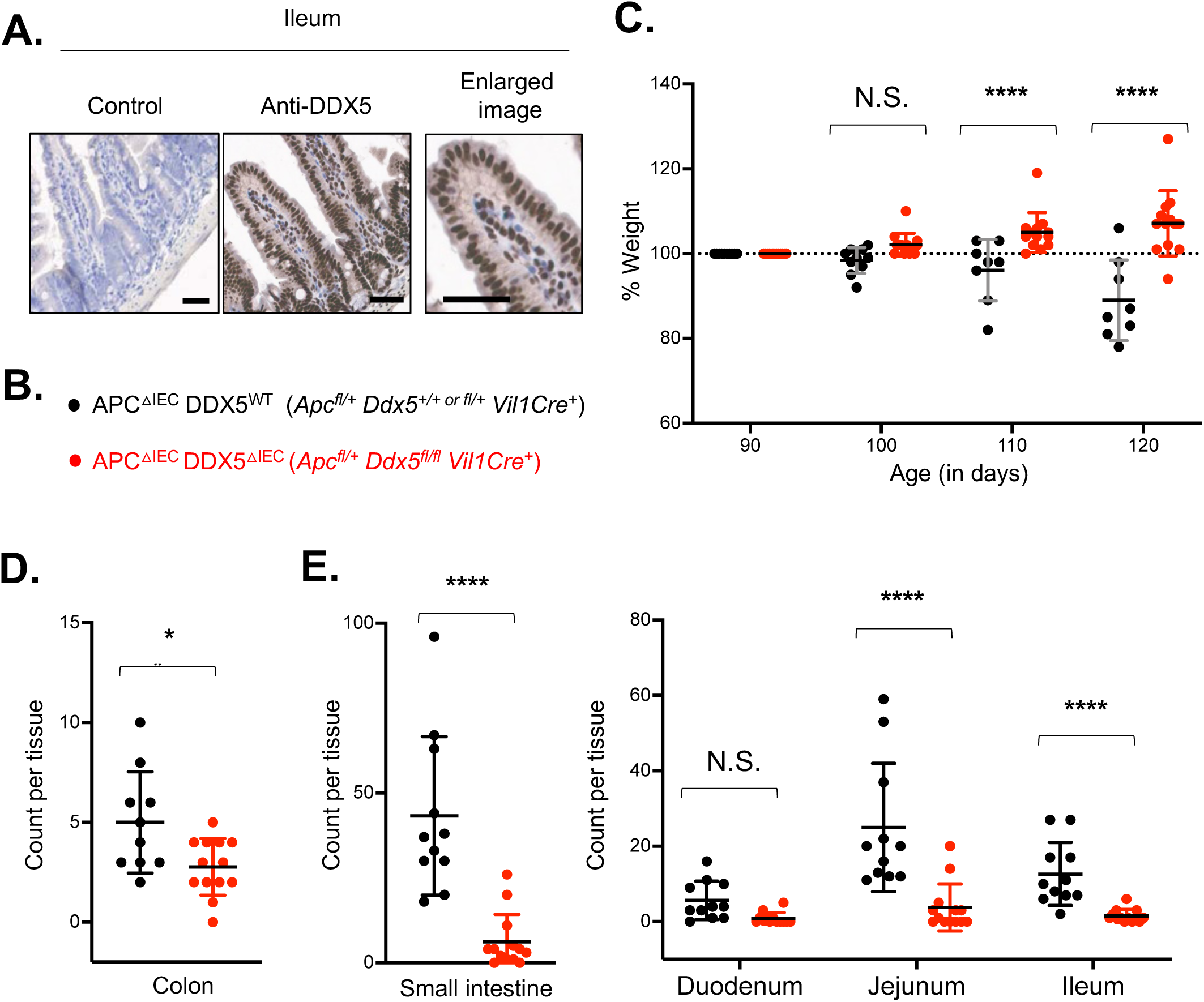
DDX5 also promotes tumorigenesis in the small intestine. A. Representative images from IHC analysis of DDX5 in the ileum of WT mice. Enlarged image is shown on the right. Scale bar represents 50 μm. B. Genotypes of tumor bearing APC^ΔIEC^DDX5^WT^ and APC^ΔIEC^ DDX5^ΔcIEC^ littermates used in C-E. C. Weight changes in DDX5 sufficient and DDX5 knockout tumor-bearing mice from day 90 to day 120 post-birth. Each dot represents one mouse. Data shown are means ± standard deviations. N.S. not significant. **** p<0.0001 (multiple *t*-test). D. Macroscopic tumor counts in the colon. Each dot represents one mouse. Counts from DDX5 sufficient samples are shown in black (n=12) and counts from DDX5 knockouts are shown in red (n=13). Data shown are means ± standard deviations. * p<0.05 (*t*-test). E. Left: Total macroscopic tumor counts in the small intestine. Right: Macroscopic tumor counts in different segments of the small intestine. Each dot represents one mouse. Counts from DDX5 sufficient samples are shown in black (n=12) and counts from DDX5 knockouts are shown in red (n=13). Data shown are means ± standard deviations. N.S. not significant. **** p<0.0001 (multiple *t*-test).

Unlike IECs in the colon, small intestine IECs expressed limited *C3* (Fig. 1G) and the expressions of β-Catenin target genes there were DDX5-independent (Fig. S8A). Hence, we speculated that DDX5 likely target other small intestine-specific RNA(s) involved in promoting tumorigenesis. Small intestine transcriptome analyses revealed DDX5-dependent programs that were distinct from those observed in the colon (Fig. 5A). The 314 DDX5-dependent small intestine IEC transcripts were enriched in pathways involved in immune responses and lipid metabolism (Fig. 5B). In particular, fatty acid-binding protein (FABP1) mRNA and protein were significantly reduced in DDX5-deficient small intestine IECs (Fig. 5C-D and Fig. S9A). While *C3* was selectively expressed in the colon, *Fabp1* was uniquely found in small intestine IECs (Fig. 5E), consistent with previous reports [56, 57]. Importantly, FABP1 knockout mice phenocopies *Ddx5*^ΔIEC^ mice and are protected from small intestine tumorigenesis [11]. These results reveal that DDX5 to be a critical upstream regulator of FABP1 and tumorigenesis in the small intestine.

**Figure 5.**
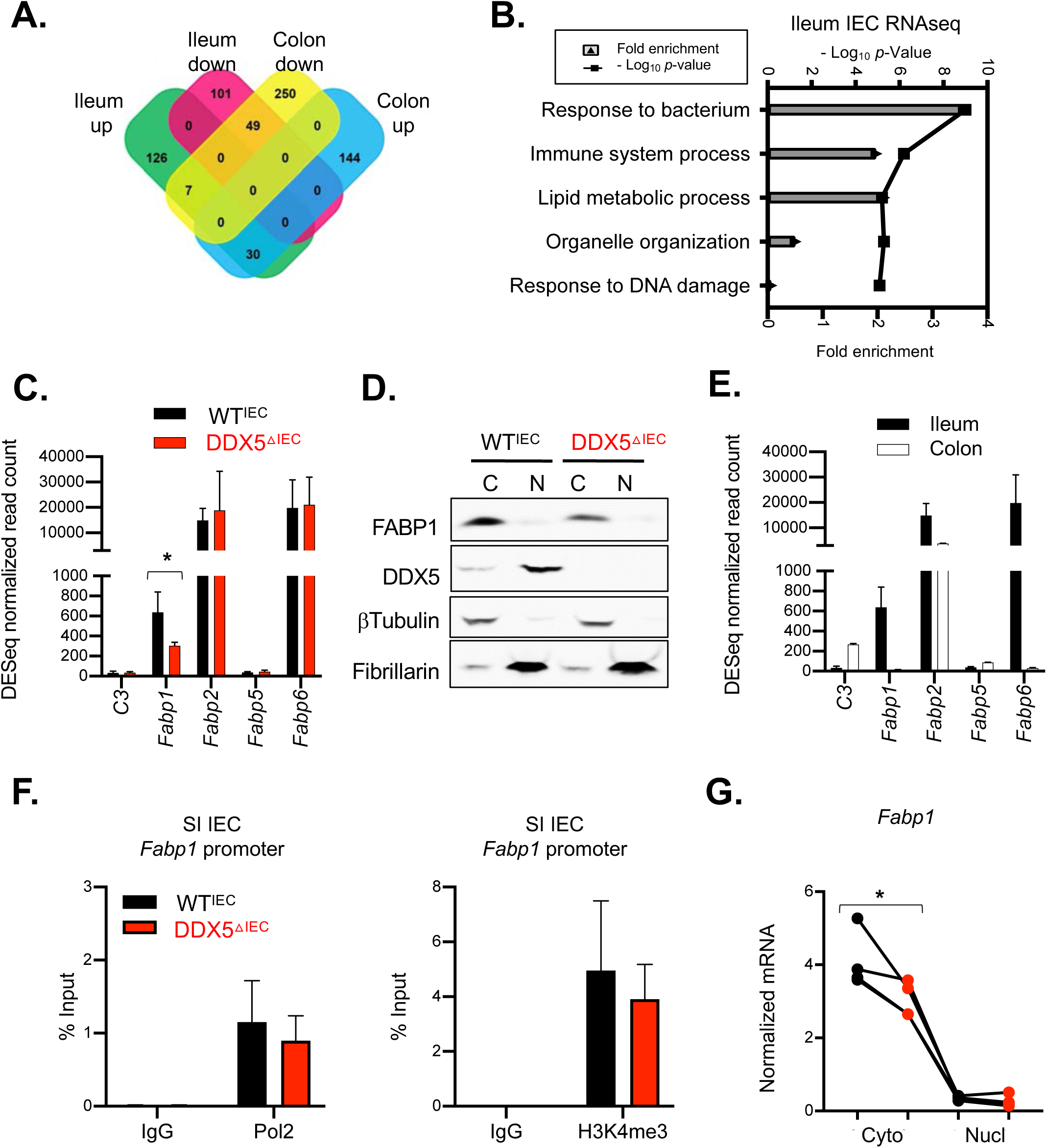
DDX5 regulates overlapping and distinct RNA programs in the small intestine and colon. A. Venn diagram of the overlapping and distinct DDX5-dependent transcripts from the small intestine and colon defined as Log_2_ fold change of ≥0.5 or ≤-0.5 and *p*-Value <0.05. B. The top 5 enriched Gene Ontology pathways identified from ileal transcripts differentially expressed in WT and DDX5^ΔIEC or ΔT^ cells defined as Log_2_ fold changes of ≥ 0.5 or ≤-0.5 and *p*-Values <0.05. All expressed genes (RNAseq count greater than 10) in the corresponding cell type were used as background. C. Normalized RNAseq read counts of transcripts encoding members of the FABP family in ileal IECs from WT^IEC^ and DDX5-deficient mice. * p<0.05 (DEseq). D. Representative western blots for FABP1, DDX5, βTubulin, and Fibrillarin in cytoplasmic (C) and nuclear (N) extracts of small intestine IECs from WT and DDX5^ΔIEC^ mice. Experiments were repeated three times using independent biological samples with similar results. E. Normalized RNAseq read counts of transcripts encoding members of the FABP family in the ileum and colon of steady-state WT mice. F. Chromatin immunoprecipitation assay using RNA polymerase II and H3K4me3 antibodies in ileal IECs from 2 independent pairs of WT^IEC^ and DDX5-deficient mice. G. RNAs from the nuclear and cytoplasmic fractions of small intestine IECs harvested from WT and DDX5^ΔIEC^ mice were evaluated by qPCR for *Fabp1*. This experiment was repeated on three pairs of independent samples. * p<0.05 (*t*-test).

ChIP assays of H3K4me3 and RNA polymerase II suggested that a similar level were found at the *Fabp1* gene promoter in small intestine IECs from WT^IEC^ and DDX5^ΔIEC^ littermates (Fig. 5F). In addition, similar abundance of mature *Fabp1* RNAs were found in the nucleus of WT^IEC^ and DDX5^ΔIEC^ IECs, but cytoplasmic mature *Fabp1* RNAs were significantly lowered in the DDX5^ΔIEC^ IECs (Fig. 5G). Together, these results point to the likelihood of DDX5 regulating small intestine *Fabp1* expression at a post-transcription step, similar to what we observed for *C3* in colonic IECs.

To determine whether DDX5 binds to and regulates *Fabp1* transcripts directly, eCLIPseq was performed using UV crosslinked small intestine IECs from WT mice (Fig. S10A). There, we found DDX5 binding to 1276 IEC RNA sites, corresponding to 466 IEC transcripts (Fig. 6A). Similar to its binding pattern in colonic IECs (Fig. 3E), DDX5 in the small intestine IECs was enriched on coding regions (Fig. 6B). On the *Fabp1* RNA, DDX5 localized to a region encoded by exon 2 (Fig. 6C). Globally, DDX5 bound coding regions of IEC transcripts were enriched with an AxGAxG motif (Fig. 6D), consistent with those identified in a previous report in human cells [58]. Similar AxGAxG motifs can be found on human *Fabp1* transcripts (Fig. 6E). Knocking down DDX5 in human epithelial Caco-2 cells also resulted in a modest decrease of human *Fabp1* expression (Fig. 6F). Together, these results demonstrate that DDX5 binding to and augmenting the expression of epithelial RNAs encoding oncogenic drivers is evolutionarily conserved (modeled in Fig. 6G).

**Figure 6.**
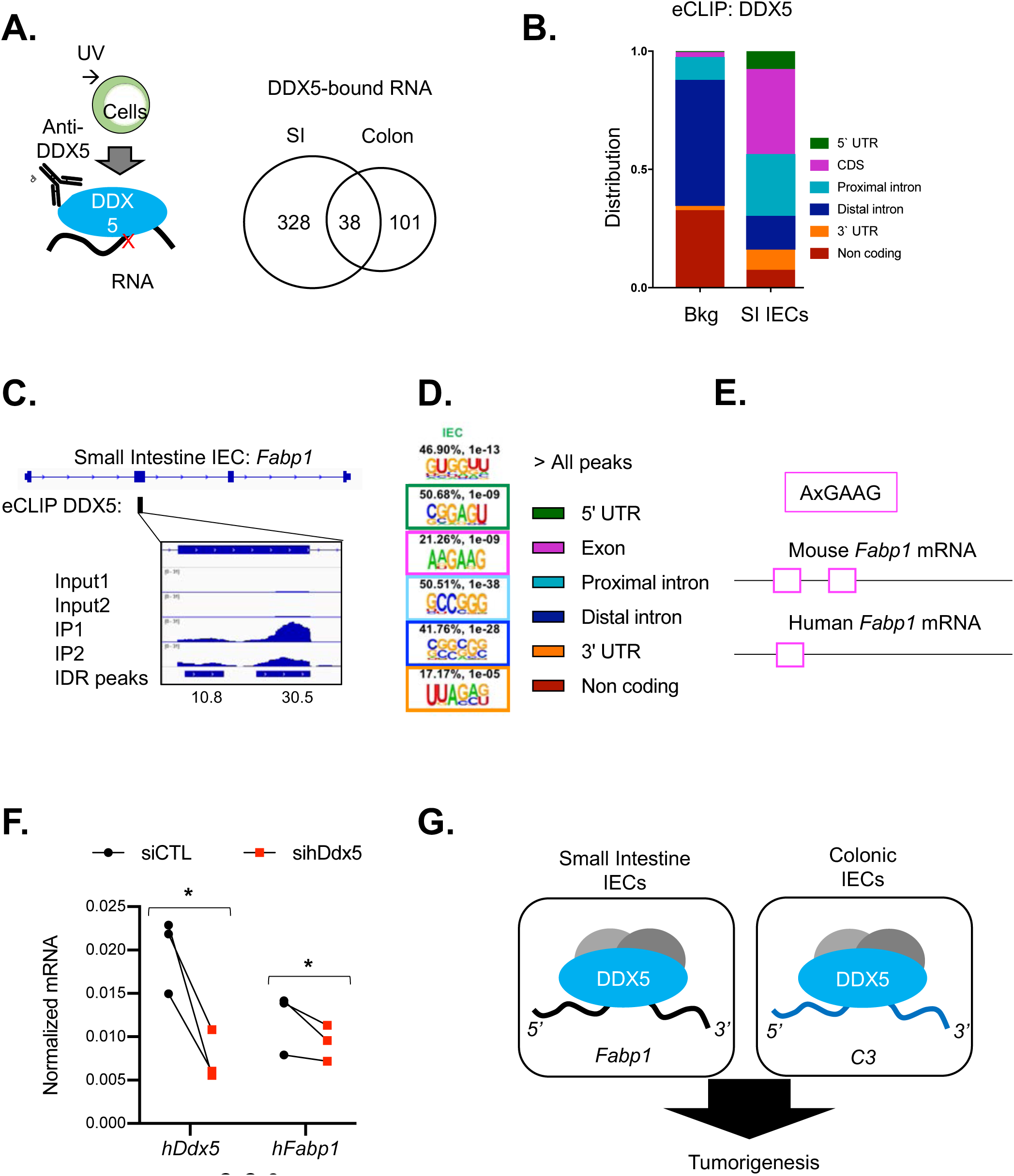
Epithelial DDX5 binds to *Fabp1* RNA and promotes its expression post-transcriptionally in the small intestine. A. Left: Schematic representation of the enhanced CLIP (eCLIPseq) experiment. Right: Venn diagram showing the overlapping and distinct DDX5-bound transcripts in the small intestine and colonic IECs. B. DDX5 binding preferences identified by eCLIPseq on different RNA regions in small intestine IECs. Background (Bkg) is defined as the RNA regions in the annotated mouse transcriptome from GENCODE. C. IGV browser displaying DDX5 binding on the *Fabp1* locus as defined by eCLIPseq. D. DDX5-bound RNA regions were enriched with unique motifs, including AxGAxG in the coding region and GC-rich motifs in non-coding regions. E. AxGAxG motif localization on the mouse and human *Fabp1* transcripts. F. RNAi mediated knockdown of human DDX5 in Caco-2 cells. Expression of human *Ddx5* and *Fabp1* was assessed 48hrs post-transfection by qPCR. Results were averaged of three independent replicates. * p<0.05 (*t*-test). G. Working model: DDX5 targets tissue-specific RNAs to promote intestine tumorigenesis

## Discussion

Intestinal cancer is the fourth most deadly cancers worldwide [59]. In these cancers, DDX5 is often mutated or overexpressed [60]. The abnormally high expression of DDX5 predicts poor patient survival [26-28]. Previous studies in cultured cancer cell lines suggest that DDX5 promotes oncogene expressions downstream of β-catenin [41, 42]. Here, our *in vivo* studies revealed an unappreciated role of DDX5 in promoting the expression of complement protein C3, a potent inducer of the Wnt/β-catenin cascade that drives tumorigenesis in the colon [44].

In the small intestine, however, we showed that DDX5 promotes tumorigenesis by targeting fatty acid-binding protein, FABP1, instead. Altered fatty acid metabolism is one hallmark of cancer [61]. Highly proliferative cells require large amounts of fatty acid building blocks from exogenous sources and/or *de novo* synthesis to sustain the building of cell membranes and organelles. For example, previous reports suggest that the upregulation of acyl-CoA synthetase ACSL4 promotes tumor cell survival in human colon adenocarcinomas [62], and that fatty acid-binding proteins can channel lipids from surrounding tissues to fuel further tumor growth [63]. Future metabolomic studies will be needed to further elucidate the DDX5-FABP1 dependent lipidome in the small intestine IECs.

Mechanistically, our *in vivo* characterization of the DDX5 RNA interactome and regulome revealed several surprises. Previous report using the human myelogenous leukemia cell line (K562) suggests that DDX5 binding on RNAs is preferentially localized to introns and 5’ UTRs [31]. Binding of DDX5 to introns of pre-mRNAs can facilitate transcript splicing and regulate protein translation [31-33, 64, 65]. In mouse IECs, we demonstrated that DDX5 preferentially localized to coding regions of RNAs. These results suggest that DDX5 binding to RNAs is likely tissue and cell-type specific. Future studies to uncover DDX5 protein interactome in different lineages will be helpful to uncover the molecular basis underlying how such binding specificities can be achieved. Importantly, we have also identified a large number of DDX5-bound epithelial RNAs that showed little dependence on DDX5 at the RNA level. It remains to be investigated whether DDX5 contributes to their structure, trafficking, and/or protein translation properties.

In summary, we report that epithelial DDX5 targeting of tissue-specific RNAs drives tumorigenesis in different parts of the intestine. These findings present DDX5 as a central regulator of oncogene expression and clinical potential for targeting DDX5 to treat intestinal cancers.

## Supplementary Figures

**Figure S1.**
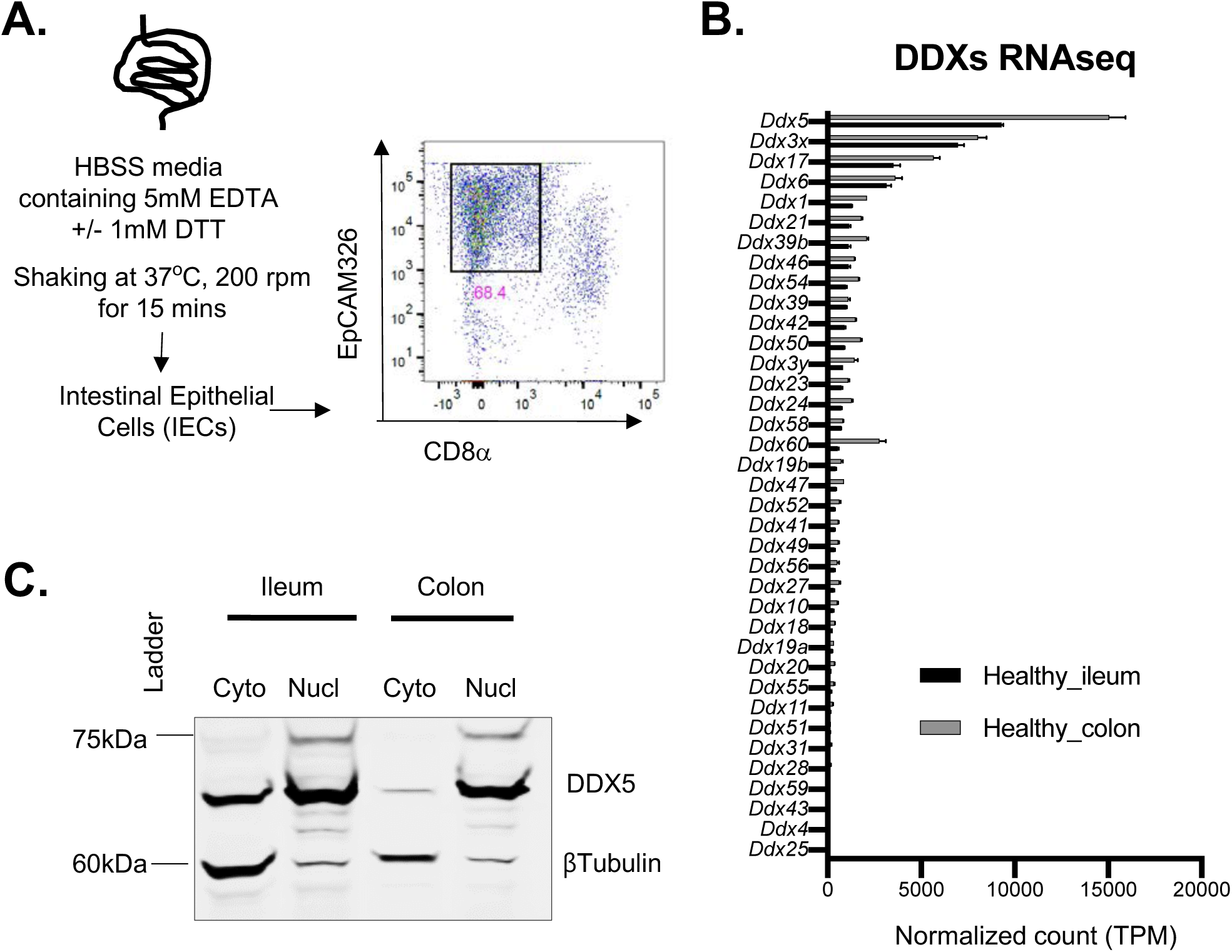
Expression of DDXs in IECs. A. Workflow for harvesting IECs from the intestine. Flow analyses confirmed EpCAM (CD326)-expressing IECs were enriched following EDTA fractionation of the small intestine. Gated on live singlets cells. B. RNA expression of DDXs in ileal and colonic IECs. C. Representative western blot of WT IEC lysates showing that DDX5 was present in the nucleus and cytoplasm. Blots were also probed with βTubulin to confirm proper nuclear and cytoplasmic fractionation.

**Figure S2.**
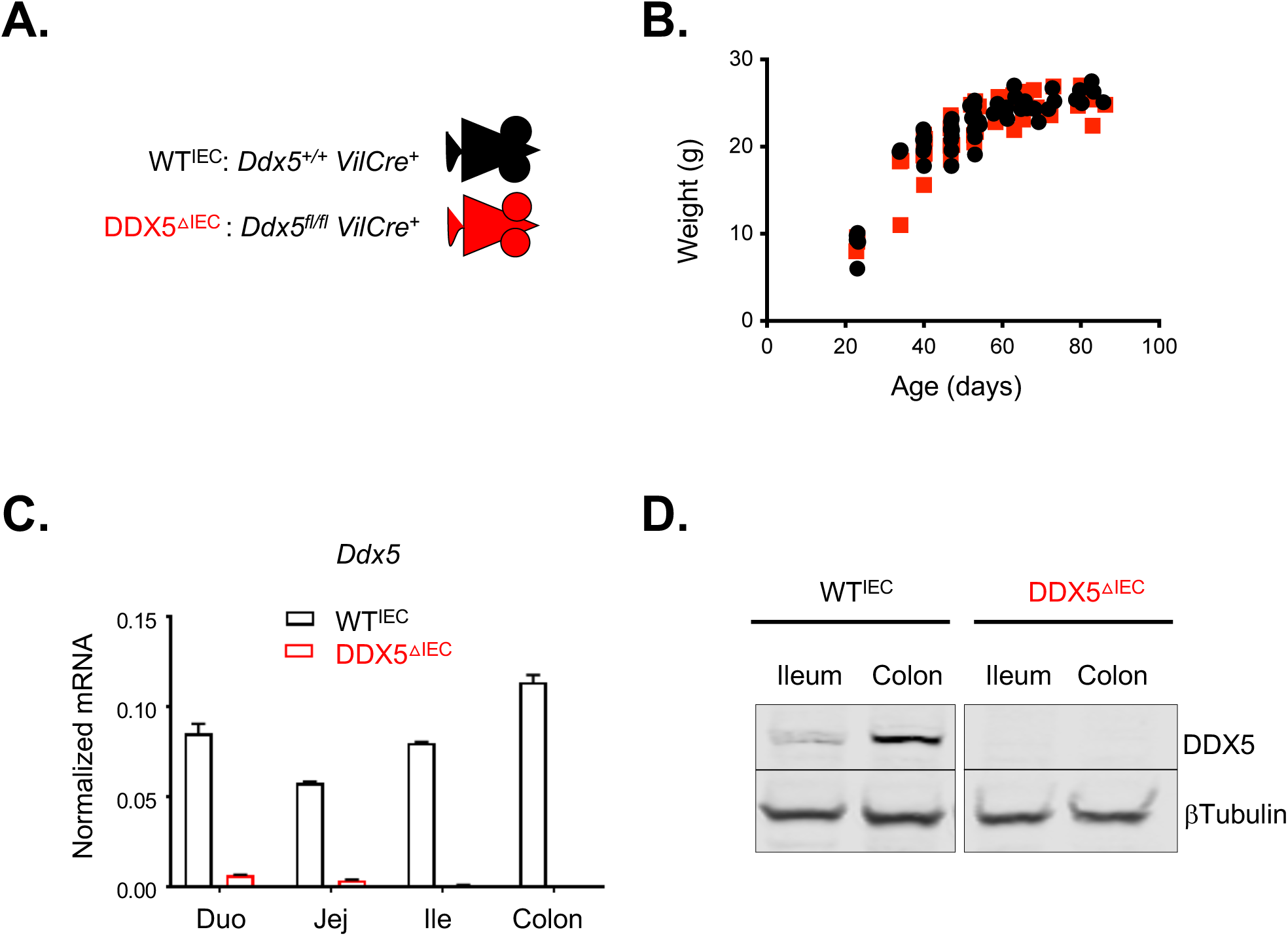
Generation of the pan-epithelial DDX5 knockout mouse line. A. Genotypes of WT^IEC^ and DDX5^ΔIEC^ littermates. B. Growth curves of WT^IEC^ and DDX5^ΔIEC^ littermates. C. Representative *Ddx5* expression, as detected by qPCR, in IECs from WT^IEC^ and DDX5^ΔIEC^ littermates showing effective depletion of the target mRNA in cells from DDX5^ΔIEC^ animals. Experiments were repeated three times using independent biological samples with similar results. D. Representative western blots showing the depletion of DDX5 protein in IECs from DDX5^ΔIEC^ mice. Experiments were repeated three times using independent biological samples with similar results.

**Figure S3.**
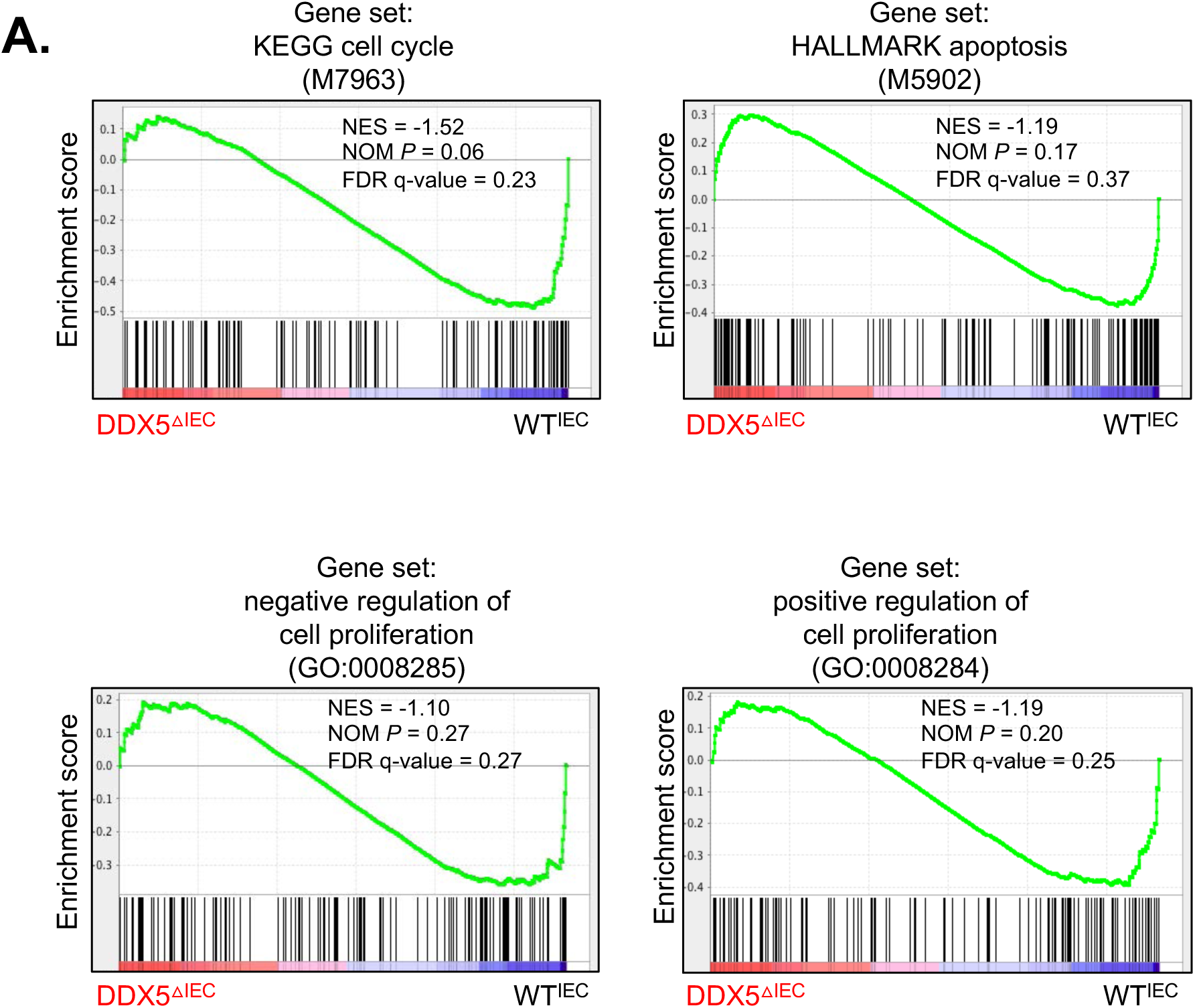
No enrichment of cell proliferation and apoptosis genes in DDX5-dependent programs from the steady-state colon. A. GSEA analysis of cell cycle, apoptosis, negative and positive regulators of cell proliferation in DDX5-deficient and DDX5-expressing colonic IECs from steady-state mice. NES, normalized enrichment score; NOM *P*, normalized *p-value*.

**Figure S4.**
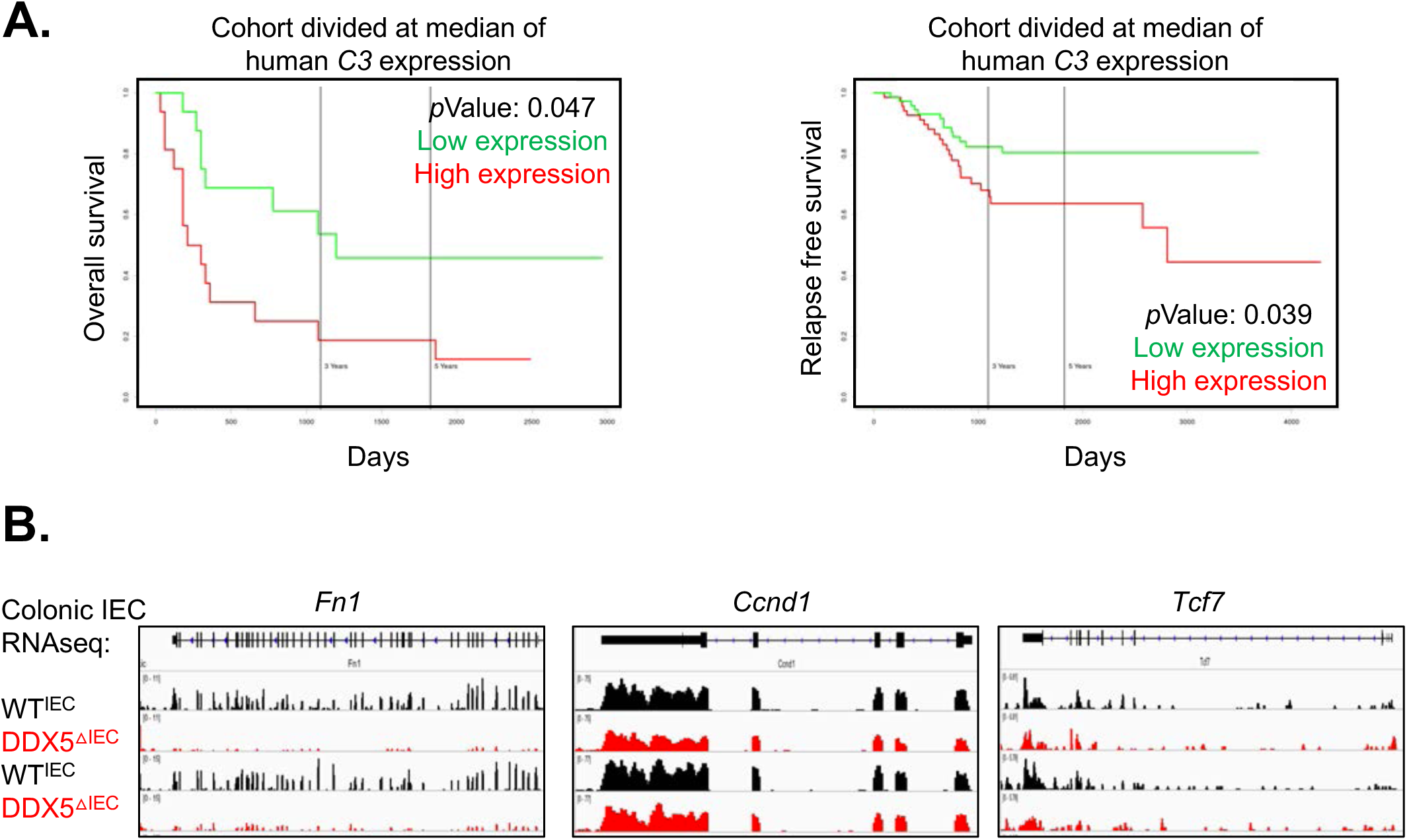
The expression of a subset of tumor stem cell-associated genes in the colon. A. PROGgeneV2 view of overall survival (left) and relapse-free survival (right) in CRC cohort (GSE16125 and GSE17536) divided at median of human *C3* gene expression. B. IGV browser displaying RNAseq signals at the *Fn1, Tcf7*, and *Ccnd1* loci.

**Figure S5.**
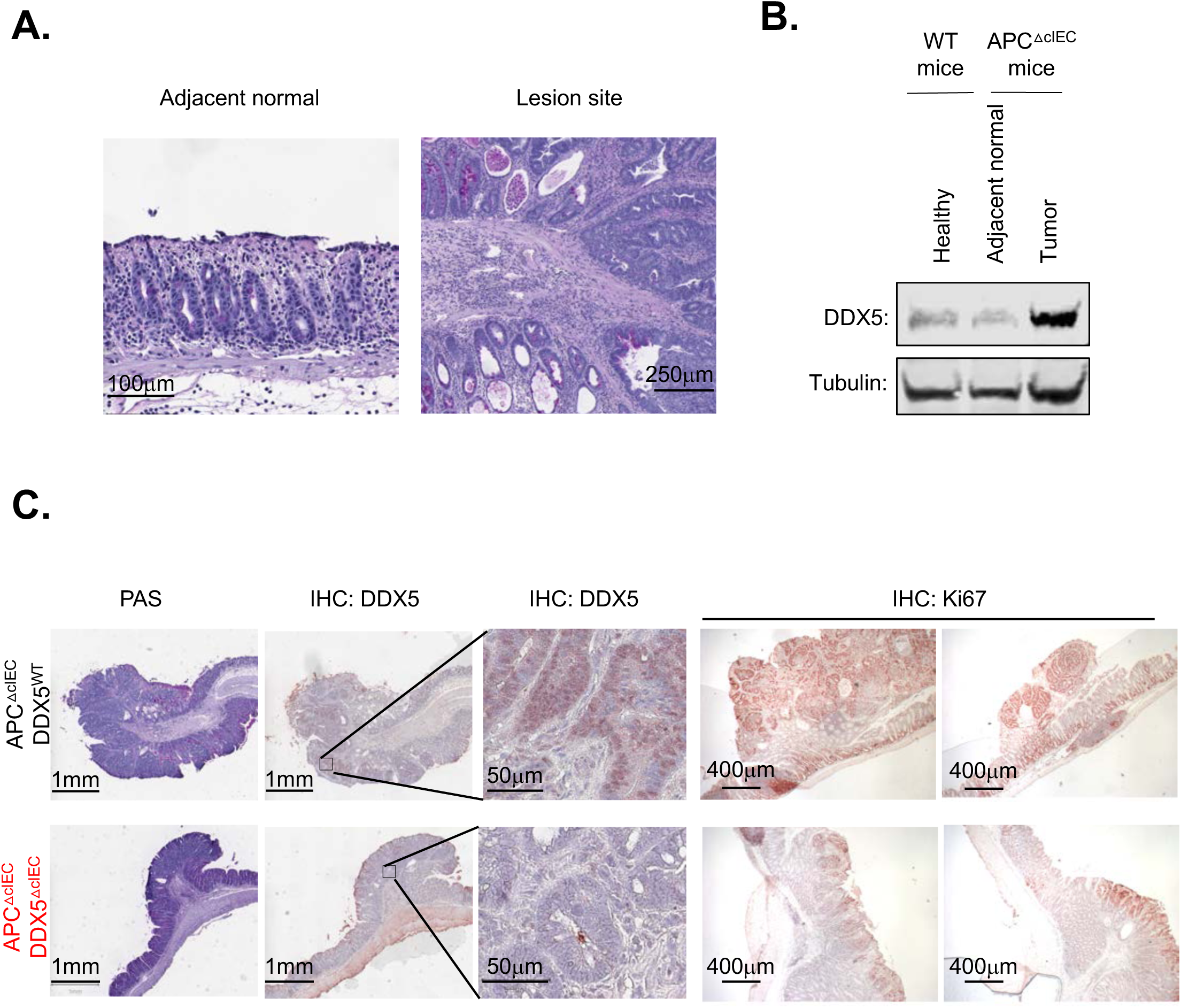
DDX5 protein expression in healthy tissues and in colonic tumors from *Apc* mutant mice. A. Representative images from PAS stained section of adjacent normal colonic epithelium and one adenoma lesion. B. Western blot analysis of DDX5 and β-Tubulin in normal and tumor tissues from the mouse colon. Experiments were repeated three times using independent biological samples with similar results. C. Representative images from PAS staining and immunohistochemical analysis of DDX5 and Ki67 in the colon of APC^ΔIEC^DDX5^WT^ and APC^ΔIEC^ DDX5^ΔcIEC^ mice.

**Figure S6.**
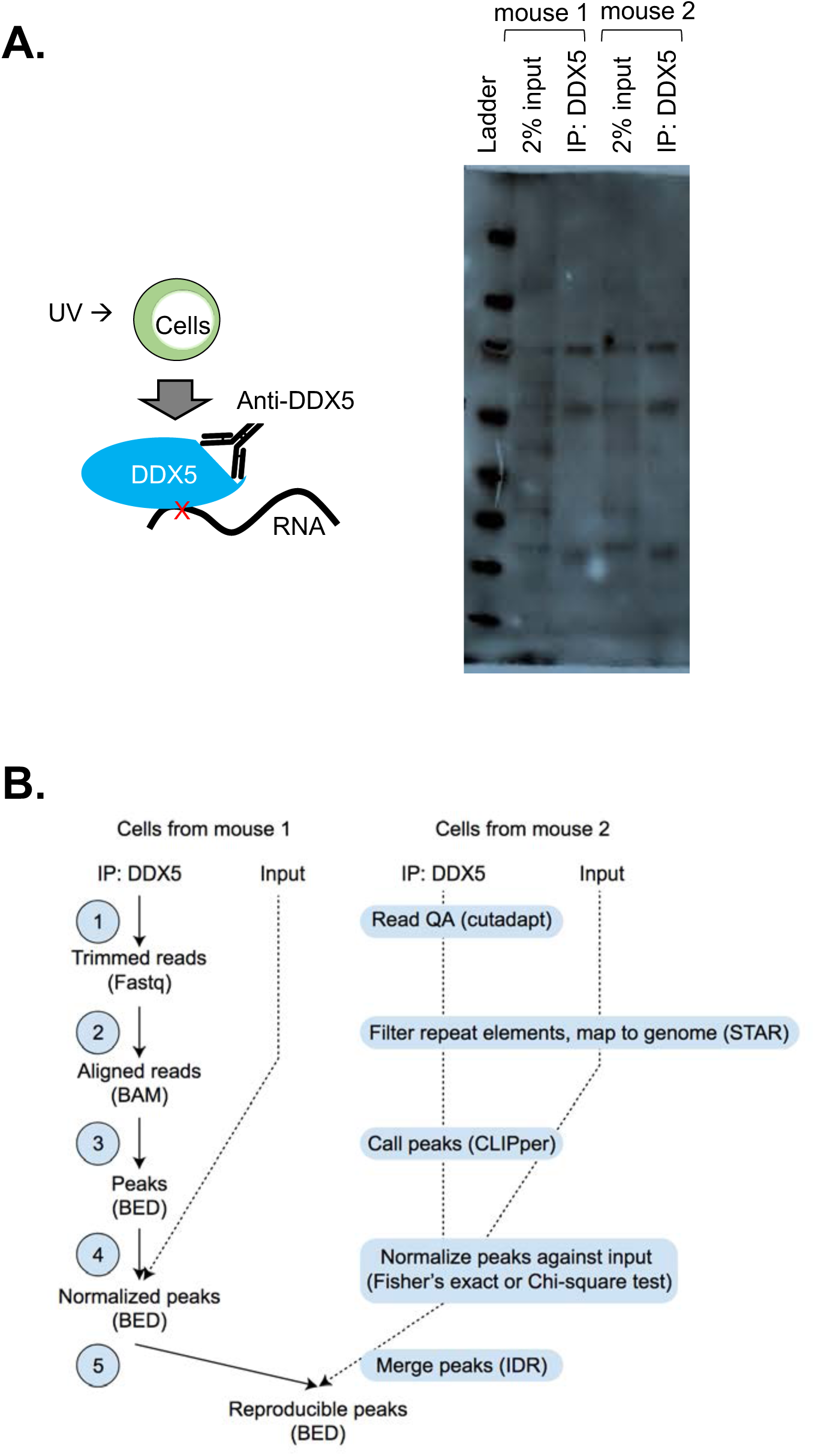
eCLIPseq analysis pipeline. A. Western confirmation of efficient immunoprecipitation of DDX5 from two independent colonic IEC lysates. B. Workflow of the eCLIPseq analysis.

**Figure S7.**
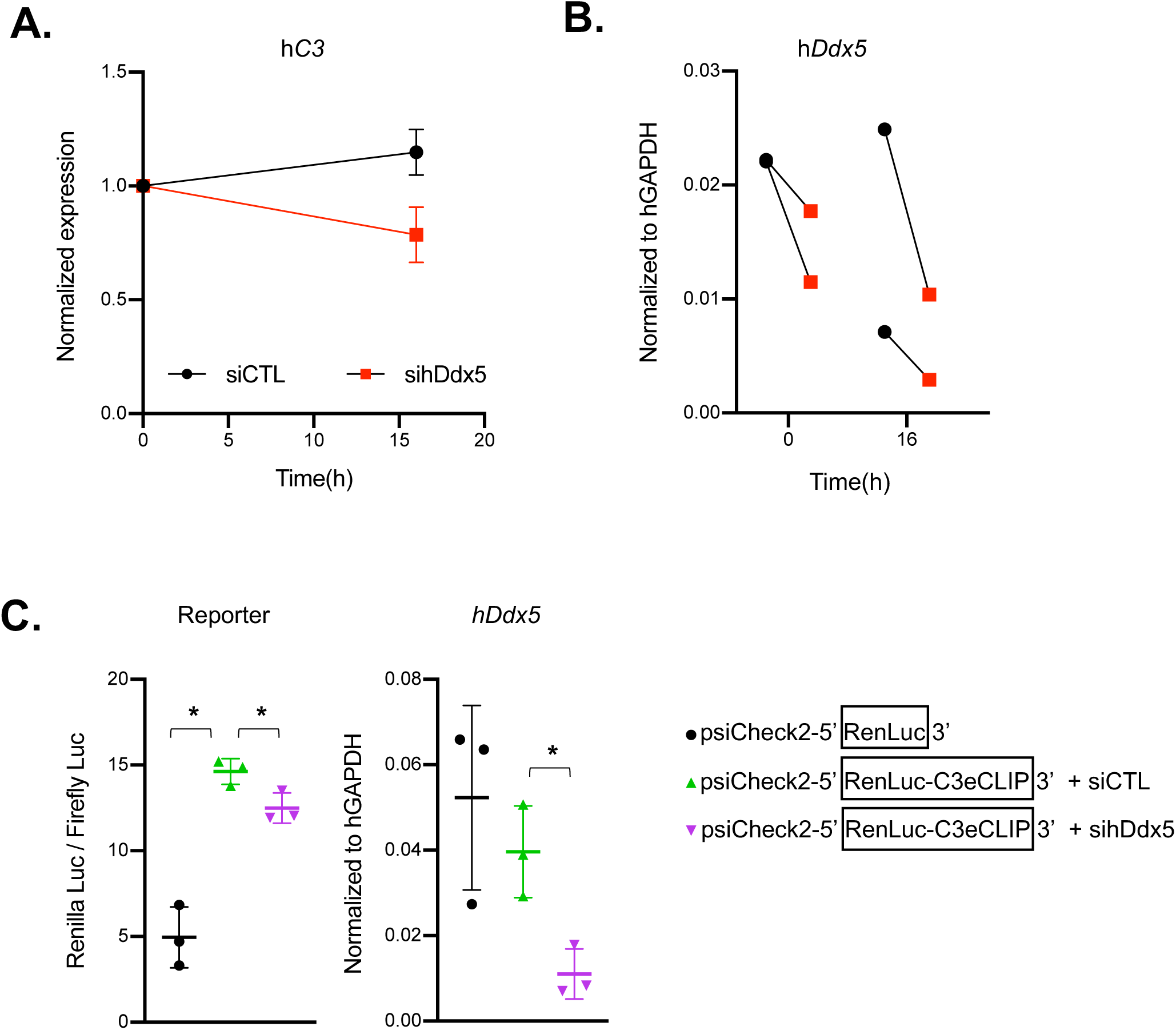
DDX5 regulates *C3* mRNA in human epithelial cells. A. RNAi mediated knockdown of human DDX5 in Caco-2 cells. Cells were transfected with control or siRNA against human *Ddx5* for 48hrs followed by 16hrs of incubation with 2μM of transcription inhibitor flavopiridol. Expressions of human *C3* were assessed by qPCR and normalized to the 0 time point. Results were averaged of two independent studies. B. Expressions of human *Ddx5* under different treatment times were assessed by qPCR and normalized with h*Gapdh*. Results were from two independent studies. C. DDX5 binding site on *C3* promotes renilla luciferase reporter activities in human SW480 cells. Results were normalized to constitutive firefly luciferase activities and averaged of three independent studies. * p<0.05 (*t*-test).

**Figure S8.**
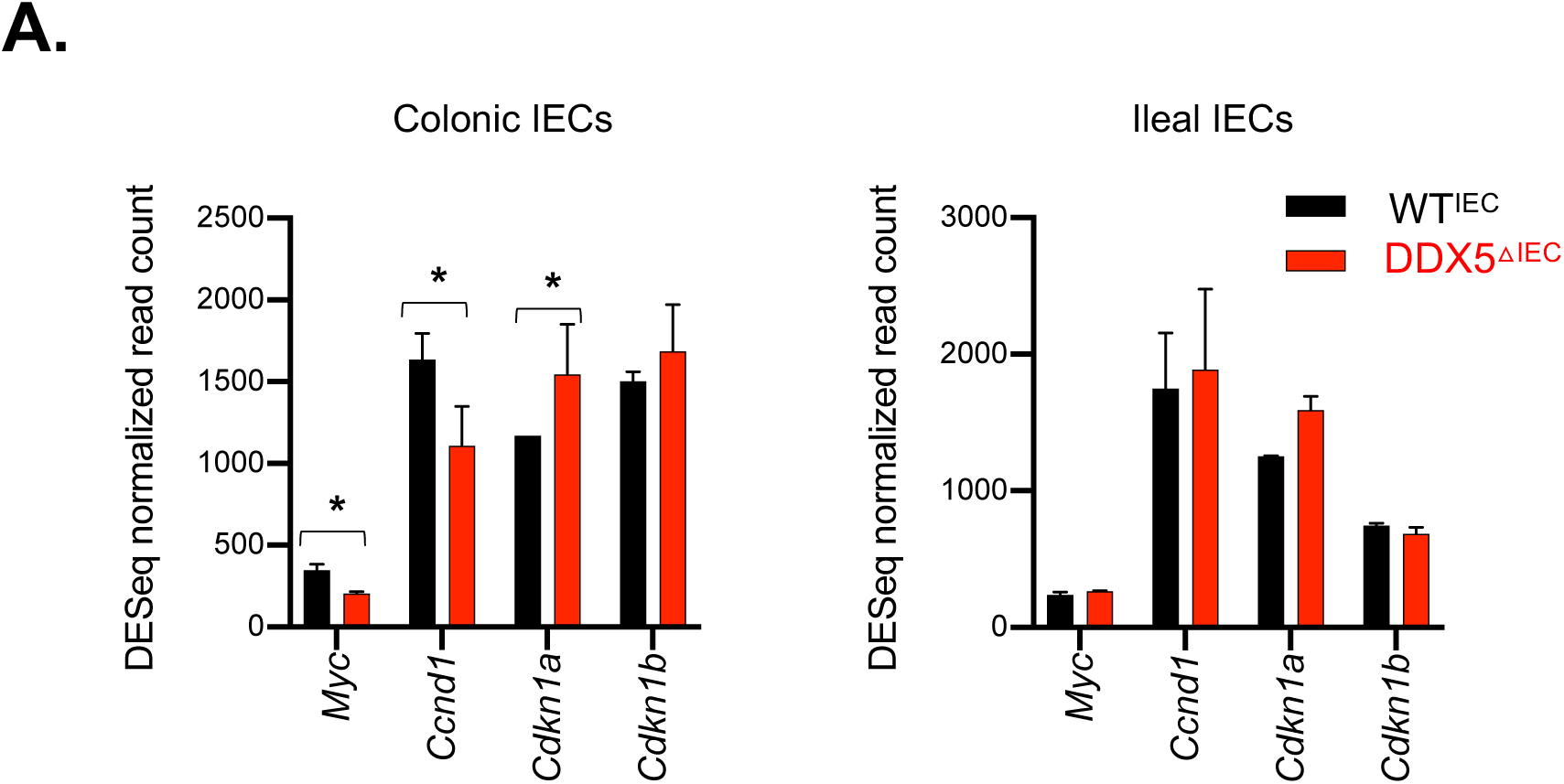
β-Catenin target gene expressions in colonic and ileal IECs. A. Normalized RNAseq read counts of known β-Catenin-regulated oncogenic transcripts in colonic and ileal IECs from WT^IEC^ and DDX5-deficient mice. * p<0.05 (DEseq).

**Figure S9.**
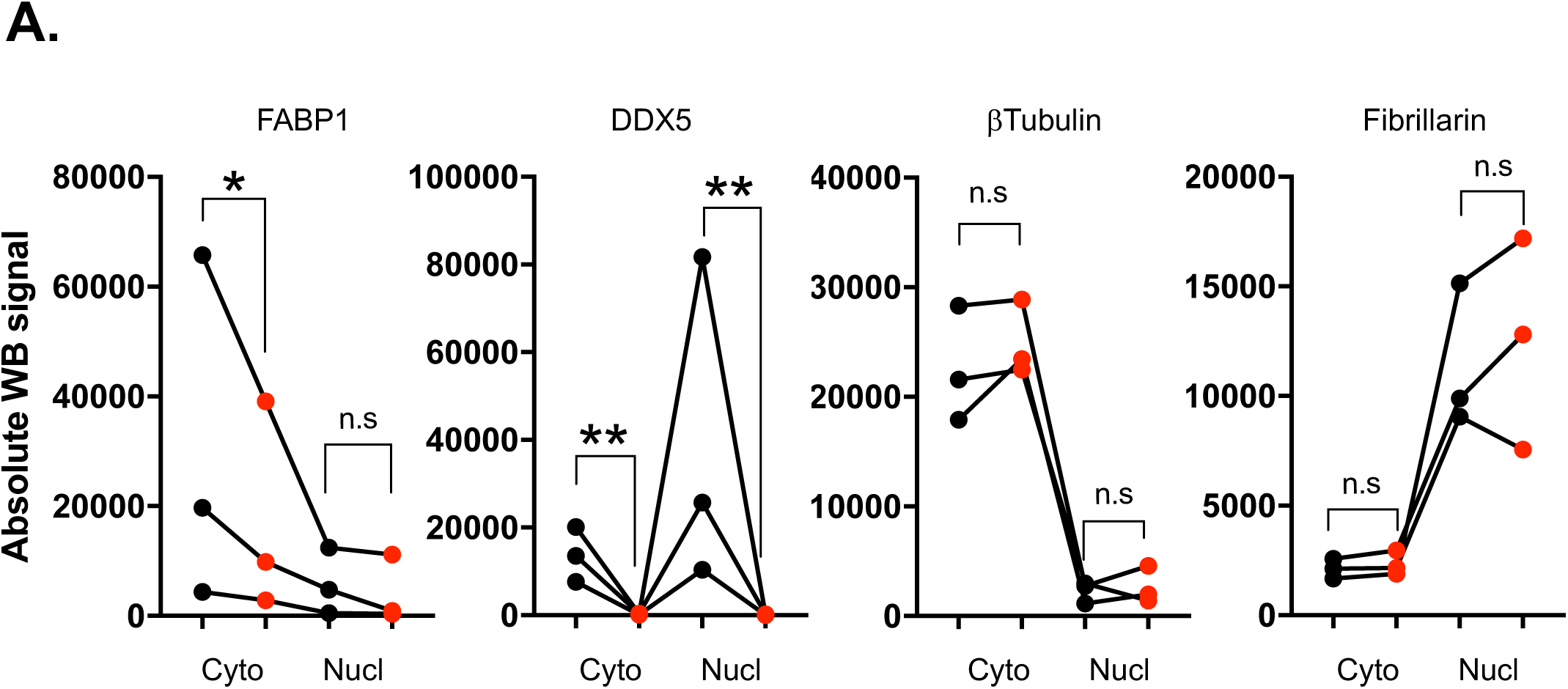
Quantification of protein expression in WT^IEC^ and DDX5^ΔIEC^ small intestine IECs. A. Quantification (LiCoR: ImageStudio) of the western blot signals from Fig. 5D (Nucl: nuclear and Cyto: cytoplasmic fractions) * p<0.05, ** p<0.01 (paired *t*-test).

**Figure S10.**
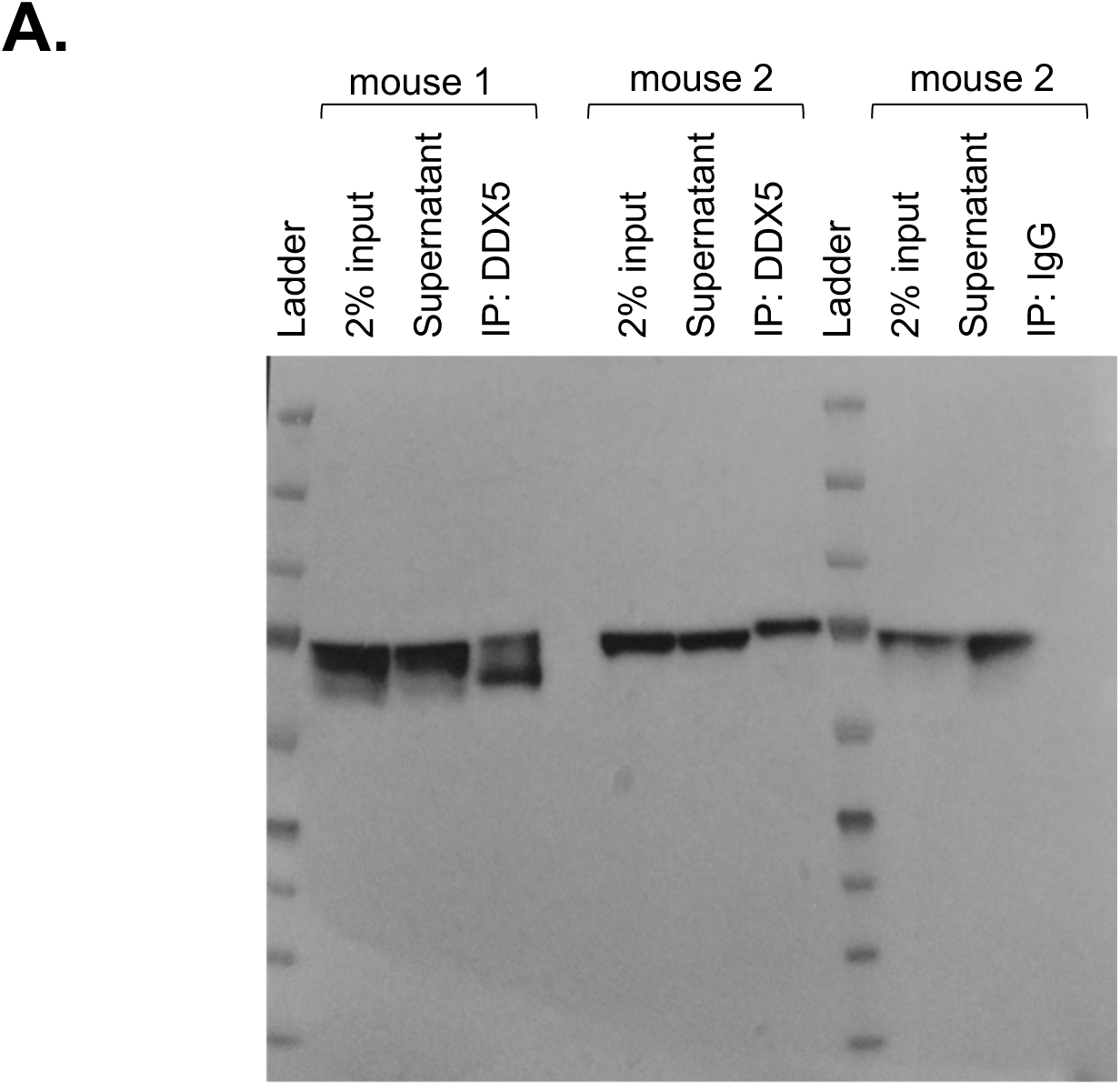
Immunoprecipitation of small intestine DDX5 for eCLIPseq analysis. A. Western confirmation of efficient immunoprecipitation of DDX5 from two independent small intestine IEC lysates.

## Methods

### Mice

C57BL/6 wild-type (Stock No: 000664) and *Villin1Cre* (Stock No: 021504) mice were obtained from The Jackson Laboratory. *Ddx5*^*flox*^ mice were obtained from Dr. Frances Fuller-Pace’s Laboratory and have been previously described in [67]. Heterozygous mice were bred to yield 6-8 week-old *Ddx5*^*+/+*^ *Villin1Cre*^*+*^ (subsequently referred to as wild-type, WT^IEC^) and *Ddx5*^*fl/fl*^ *Villin1Cre*^*+*^ (referred to as DDX5^ΔIEC^) littermates for experiments related to understanding the role of DDX5 in IECs in both the small intestine and colon. *Apc*^*flox*^ mice were obtained from Dr. Eric Fearon’s Laboratory and previously described in [51]. For our colonic tumor model, *Apc*^*fl/+*^ *Ddx5*^*+/+*^ *Cdx2Cre*^*+*^ and *Apc*^*fl/+*^ *Ddx5*^*fl/+*^ *Cdx2Cre*^*+*^ (referred as APC^ΔcIEC^ DDX5^WT^), as well as *Apc*^*fl/+*^ *Ddx5*^*fl/fl*^ *Cdx2Cre*^*+*^ (APC^ΔcIEC^ DDX5^ΔcIEC^) cohoused littermates were used. For our small intestine tumor model, *Apc*^*fl/+*^ *Ddx5*^*+/+*^ *Villin1Cre*^*+*^ and *Apc*^*fl/+*^ *Ddx5*^*fl/+*^ *Villin1Cre*^*+*^ (referred as APC^ΔIEC^ DDX5^WT^), as well as *Apc*^*fl/+*^ *Ddx5*^*fl/fl*^ *Villin1Cre*^*+*^ (APC^ΔIEC^ DDX5^ΔIEC^) cohoused littermates were used. All animal studies were approved and followed the Institutional Animal Care and Use Guidelines of the University of California San Diego.

### Epithelial cell harvest

Steady-state intestinal epithelial and lamina propria cells were harvested as previously described [68]. Briefly, after removing mesenteric fat and Peyer’s patches, the proximal 1/3, middle 1/3, and distal 1/3 of the small intestine were designated as duodenum, jejunum, and ileum, respectively. To isolate intestinal epithelial cells (IECs), intestine tissues were first incubated in 5 mM EDTA in HBSS containing 1 mM DTT for 20 min at 37°C with shaking, and then incubated in a second wash of 5 mM EDTA in HBSS without DTT for 20 min at 37°C with agitation. Suspended cells from the EDTA washes were pooled as “IECs”. Colons were processed similarly.

### Intestine organoid cultures

Isolated colonic crypts were embedded in Corning® Matrigel® Matrix Corning™ Matrigel™ GFR Membrane Matrix (Fisher Scientific CB40230C) and seeded onto pre-warmed 24-well plates (CytoOne) and overlaid with conditioned media as described in [66]. The organoid images were acquired using fluorescence microscopy (Fisher Scientific 11350119).

### Histology and IHC

Ileal and colonic tissues were fixed overnight in 10% formalin at room temperature. Paraffin-embedded tissues were sectioned into 5 μm slices, stained with H&E, PAS, or IHC (see Table S1 for antibody information). Briefly, paraffin sections were de-paraffinized and rehydrated with TBST washes between each step (Tris-buffered saline pH 7.8 with 0.1% Tween-20). Sections were blocked first against endogenous peroxidases (immersed for 30 minutes in 0.3% H_2_O_2_), and then blocked against endogenous biotin using unlabeled streptavidin) and excess free biotin. Antigen retrieval was induced by heating the slide for 5 minutes twice in 10 mM sodium citrate buffer pH 6.0, followed by 20 minutes of cooling. Finally, sections were blocked against non-specific hydrophobic interactions with 1% BSA/TBST (bovine serum albumin). Staining was then performed with either the negative control IgG antibody or anti-DDX5 and anti-Ki67 (1:100) antibodies overnight in a humid chamber at 4 °C. The next day, sections were washed with TBST and then sequentially overlaid with biotinylated goat anti-rabbit (Jackson ImmunoResearch, 111-065-045) at 1:500, followed by HRP-labeled Streptavidin (Jackson ImmunoResearch, 16-030-084) at 1:500. Substrate was then overlaid (AEC from Vector labs following directions) for 30 minutes followed by nuclear counterstain with Mayer’s hematoxylin. Images were acquired using the AT2 Aperio Scan Scope (UCSD Moores Cancer Center Histology Core).

### Western blot

For whole cell lysates, cells were lysed in 25 mM Tris pH 8.0, 100 mM NaCl, 0.5% NP40 with protease inhibitors for 30 min on ice. Samples were spun down at 14,000 x *g* for 15 min, and soluble protein lysates were harvested. The NE-PER™ kit (ThermoFisher Scientific) was used for cytoplasmic and nuclear fractionation studies. 30-50μg protein were loaded on each lane. Blots were blocked in Odyssey Blocking buffer (Li-CoR) and probed for the desired proteins. Following incubation with respective IRDye secondary antibody (Li-CoR^®^), infrared signals on each blot were measured on the Li-CoR Odyssey CLX. The primary antibodies used in this study are listed in Table S1.

### cDNA synthesis and qPCR

Total RNA was extracted with the RNeasy Plus kit (QIAGEN) and reverse transcribed using iScriptΪ™ Select cDNA Synthesis Kit (Bio-Rad). Real time RT-PCR was performed using iTaq™ Universal SYBR^®^ Green Supermix (Bio-Rad). For IECs and tumor RNA expression, data was normalized to *Gapdh*. For organoid RNA expression, data were normalized to *Actβ*. Primers were designed using Primer-BLAST to span across splice junctions, resulting in PCR amplicons that span at least one intron. Primer sequences are listed in Table S2.

### RNAseq

Ribosome-depleted RNAs were used to prepare sequencing libraries. 100 bp paired-end sequencing was performed on an Illumina HiSeq4000 by the Institute of Genomic Medicine (IGM) at the University of California San Diego. Each sample yielded approximately 30-40 million reads. Paired-end reads were aligned to the mouse mm10 genome with the STAR aligner version 2.6.1a [69] using the parameters: “-- outFilterMultimapNmax 20 --alignSJoverhangMin 8 --alignSJDBoverhangMin 1 -- outFilterMismatchNmax 999 --outFilterMismatchNoverReadLmax 0.04 --alignIntronMin 20 --alignIntronMax 1000000 --alignMatesGapMax 1000000”. Uniquely mapped reads overlapping with exons were counted using featureCounts [70] for each gene in the GENCODE.vM19 annotation. Differential expression analysis was performed using DESeq2 (v1.18.1 package) [71], including a covariate in the design matrix to account for differences in harvest batch/time points. Regularized logarithm (rlog) transformation of the read counts of each gene was carried out using DESeq2. Pathway analysis was performed on differentially expressed protein coding genes with minimal counts of 10, log_2_ fold change cutoffs of ≥ 0.5 or ≤-0.5, and *p*-Values <0.05 using Gene Ontology (http://www.geneontology.org/) where all expressed genes in the specific cell type were set as background.

Gene set enrichment analysis was carried out using the pre-ranked mode of the GSEA software with default settings [72, 73]. The gene list from DEseq2 was ranked by calculating a rank score of each gene as −Log_10_(*p-value*) × sign (Log_2_ FoldChange), in which FoldChange is the fold change of expression in DDX5^ΔIEC^ over those found in WT^IEC^.

### <u>E</u>nhanced <u>C</u>ross-<u>L</u>inked <u>I</u>mmuno<u>p</u>recipitation (eCLIPseq)

eCLIPseq analysis was performed as previously described [55]. For IEC eCLIPseq, cells were isolated from two 8-10 week-old wild-type (C57BL/6) female mice, as described above, and 50 million cells from each mouse were used in the two biological replicates. The cells were subjected to UV-mediated crosslinking, lysis, and treatment with limiting amounts of RNases, followed by immunoprecipitation (IP) of the DDX5-containing RNA complexes. RNA fragments protected from RNase digestion were subjected to RNA linker ligation, reverse-transcription, and DNA linker ligation to generate eCLIPseq libraries for high-throughput Illumina sequencing.

Peak regions were defined using CLIPper first on the IP sample (https://github.com/YeoLab/clipper/wiki/CLIPper-Home). Enrichment was calculated using both the IP and input samples. Log_2_ fold change was calculated as eCLIPseq reads normalized for read depth over normalized reads found at each peak region in the size-matched input sample. ENCODE Irreproducible Discovery Rate (IDR) analysis was performed on two independent biological replicates of IECs. Peaks were ranked using the entropy formula, Pi*log(Pi/Qi)/log_2_, where Pi is the probability of an eCLIPseq read at that position and Qi is the probability of input reads at that position. Results were filtered using cutoffs of 3 and 3 for log_10_ *p*-values and log_2_ fold changes, respectively, to define a set of true peaks normalized above their respective size-matched input background signal. Differentially expressed genes were defined as those with minimal counts of 5 and log_2_ fold change cutoffs of ≥ 0.4 or ≤-0.4.

### Chromatin immunoprecipitation (ChIP)

Chromatin immunoprecipitation was carried out as described previously [74]. Briefly, 20 million intestinal IECs were fixed with 1% formaldehyde at room temperature for 10 mins, and quenched with 125 mM glycine for 5 min at room temperature. All buffer compositions were described in [74]. Nuclear lysates were sonicated with a Bioruptor (Branson Sonifier Cell Disruptor 185.) at 4°C using the output setting at 4 for 10 cycles of 30 sec on and 30 seconds off. 30 μg of chromatin was used per IP. Chromatin was diluted 10X in ChIP dilution buffer supplemented with proteinase inhibitor. 5% of the total chromatin used per IP reaction was saved as input samples. 5 μg of antibody was added per 30 μg chromatin per IP reaction and incubated overnight at 4°C. The immune complexes were then incubated with 30 μL of Dynabeads™ Protein G (Thermo Fisher Scientific,10004D) for 4 hours at 4°C on rotation. After washes, protein-DNA complexes were eluted from the beads by adding 200 μL of elution buffer and incubating the beads at 65°C for 15 min with constant shaking at 1350 rpm. Eluted samples were incubated at 37°C for 30 mins with 1 μL of DNase and protease-free RNase A (10 mg/mL, Thermo Fisher Scientific EN0531). DNA and protein cross-linking was reversed by adding 8 μL of 5M NaCL and 2 μL of Proteinase K Solution (20 mg/mL, Thermo Fisher Scientific AM2546) by overnight incubation at 65C under constant shaking. Chromatin was isolated using Qiagen QIAquick PCR Purification Kit (28104) and eluted in 40 μL elution buffer. Input samples were diluted 5 times to make a 1% input control. The ChIP signals were calculated as follows: Adjusted input = Ct (Input) – 6.644. ChIP signal = 100 X POWER (2; average of adjusted Input-Ct value ChIP sample). All ChIP qPCR primers are listed in Supplementary table S1.

### RNAi in human intestinal epithelial cells

Caco-2 cell line was cultured in 1X DMEM/F12 media (Gibco, Life Technologies). The media were supplemented with 1X 10% FBS (Gibco, Life Technologies), 1mM Sodium Pyruvate (Gibco, Life Technologies) and 1% Penicillin Streptomycin (Gibco, Life Technologies). Cells were plated on a 24 well plate at 500μL/well at 2×10^5^ cells/mL one day before transfection. 100nM Human DDX5 siRNA pool (GE Healthcare Dharmacon, Catalog# D-003774-02, D-003774-03, D-003774-04, D-003774-17, see Table S3) or scramble siRNA pool (GE Healthcare Dharmacon, Catalog# D-001206-14-05) were mixed with Opti-MEM medium and Lipofectamine 3000 (L3000001, Invitrogen) reagent according to the manufacturer’s protocol. Solutions were vortexed and incubated for 5 minutes at room temperature to allow the formation of siRNA-lipid complex. 50μL of transfection/siRNA (final concentration of 50μM) mixture was added to the well and incubate at 37°C. Transcription inhibitor, flavopiridol, was purchased from Sigma (F3055). siRNA sequences are listed in Table S3. After incubation for 48 hours, the cells were treated with flavopiridol (2μM) and were collected 16 hours post treatment. RNA extraction and RT-qPCR were carried out as described above.

### Luciferase assay

SW480 cells were cultured in 1X DMEM/F12 media (Gibco, Life Technologies). 1X 10% FBS (Gibco, Life Technologies), 1mM Sodium Pyruvate (Gibco, Life Technologies) and 1% Penicillin Streptomycin were added (Gibco, Life Technologies). Cells were plated on a 96-well plate at 50 μL/well 0.5×10^5^ cells/mL one day before transfection. 100nM of siCTL or the Human DDX5 siRNA pool were mixed with Lipofectamine 3000 (L3000001, Invitrogen) in Opti-MEM medium according to the manufacturer’s protocol. Transfection mixture were incubated at room temp for 5 minutes. 10μL of transfection mixture was added to each well and incubated at 37°C for 24hrs. Next day, 1μg psiCheck2 empty vector or psiCheck2_Fabp1eCLIP plasmids were mixed with Lipofectamine 3000 (L3000001, Invitrogen) and P3000 enhancer reagent in Opti-MEM medium.10μL of transfection mixture was added to each well and incubated at 37°C. After 24 hours, the cells were harvested for dual Luciferase Assay. The psiCheck2 reporter construct containing dual *Renilla* and Firefly luciferase reporters was purchased from Promega (Promega, Madison, WI, USA). The DDX5-bound sequences identified by eCLIP IDR were cloned into a multiple cloning site located downstream of the *Renilla* translational stop codon. Both *Renilla* and Firely luciferase activities were measured using the Dual-Luciferase Reporter Assay System (Promega) according to manufacturer instructions.

### Statistical analysis

All values are presented as mean ± SD. Significant differences were evaluated using GraphPad Prism 8 software. The Student’s *t*-test was used to determine significant differences between two groups. A two-tailed *p*-value of <0.05 was considered statistically significant in all experiments.

## Author contributions

N.A. designed and performed the *in vivo* tuft cells and tumor studies. T.L. completed the human cell studies and together with I.M.S. performed the epithelial organoid assays. B.A.Y. completed the eCLIP analyses. Y.L. performed the bioinformatics analyses on the RNAseq datasets. E.M. and P.R.P. completed the double-blinded analyses of the tumor studies. N.V. completed the tumor histology studies. G.W.Y. directed the eCLIP studies and edited the manuscript. P.G. and S.D. directed the organoid studies and edited the manuscript. N.A. and T.L. wrote the manuscript with inputs from W.J.M.H.

## Acknowledgements

We thank Eric Fearon for sharing the APC conditional mutant mice previously described in [51]. We thank Frances Fuller-Pace for sharing the DDX5 conditional mutant mice previously described in [75]. We thank Giuseppe Di Caro for Apc mouse colony breeding advice. We thank Jean Y.J. Wang for critical reading of this manuscript.

N.A., T.L., Y.L., E.M., and W.J.M.H. are partially funded by the Edward Mallinckrodt, Jr. Foundation and the National Institutes of Health (NIGMS_1R01 GM124494). RNAseq was conducted at the IGM Genomics Center, University of California San Diego, with support from MCC grant # P30CA023100. We thank the UC San Diego HUMANOID Core and the Moores Cancer Center Histology Core Tissue Technology Shared Resource for the IHC analysis, supported by the National Cancer Institute Cancer Center Support Grant (CCSG Grant P30CA23100). The authors declare no competing financial interests.

---

**Table S1.** Antibody information.

| **Name** | **Catalog number** | **Company** | **Assay** |
| --- | --- | --- | --- |
| DDX5 | ab53216 | Abcam | eCLIP |
| DDX5 | ab10261 | Abcam | Western |
| DDX5 | 9877S | Cell Signaling | IHC |
| FABP1 | 13368T | Cell Signaling | Western |
| β-Tubulin | 2128S | Cell Signaling | Western |
| Fibrillarin | 2639S | Cell Signaling | Western |
| Normal Rabbit IgG | 12-370 | MILLIPORE | ChIP |
| RNA Polymerase II | GTX102535 | GeneTex | ChIP |
| H3K4me3 | ab8580 | Abcam | ChIP |

**Table S2.**
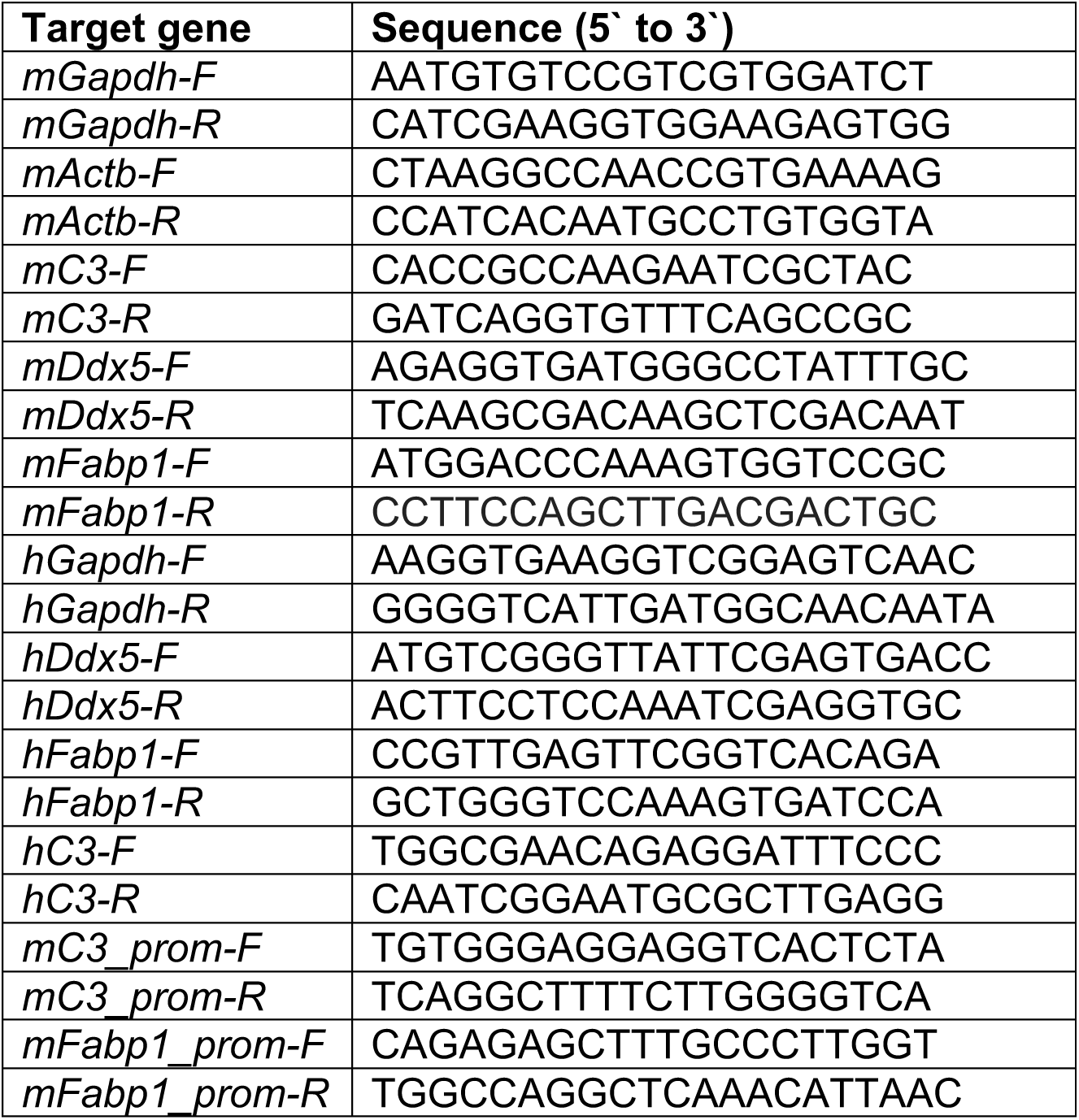
Primer information.

**Table S3.**
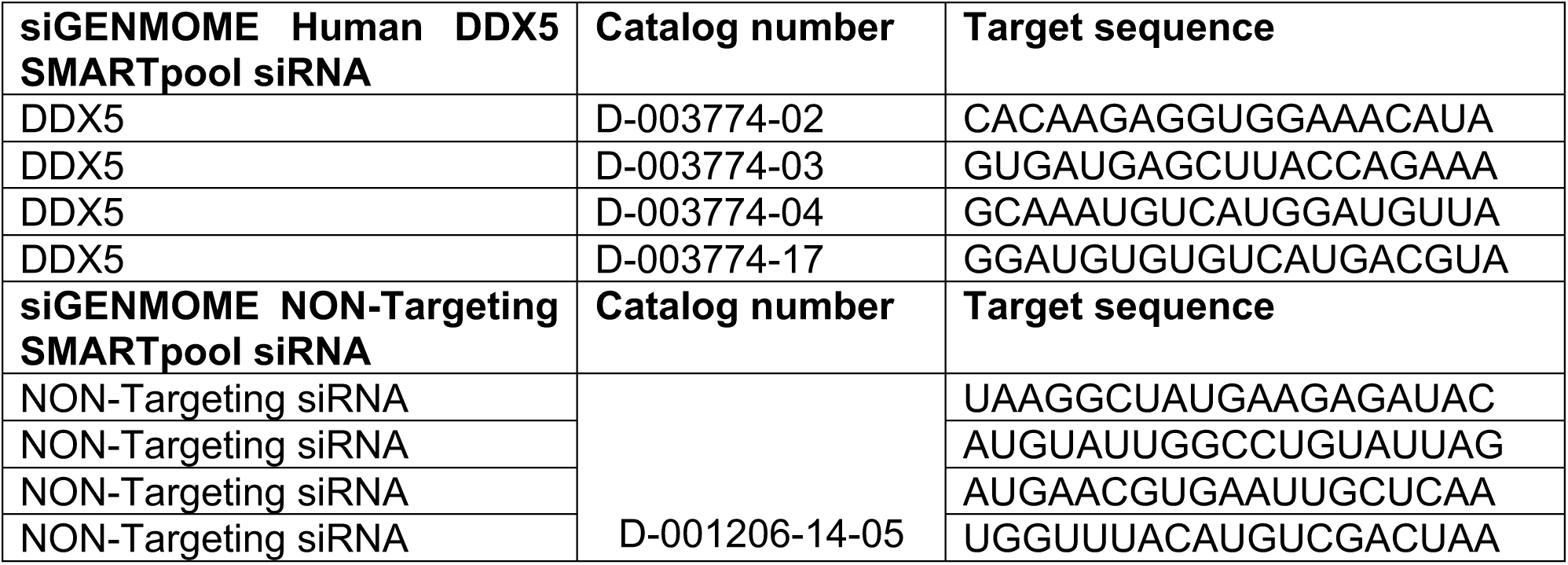
siRNA information:

